# A network information theoretic framework to characterise muscle synergies in space and time

**DOI:** 10.1101/2021.10.15.464450

**Authors:** David Ó’ Reilly, Ioannis Delis

## Abstract

**Objective:** Current approaches to muscle synergy extraction rely on linear dimensionality reduction algorithms that make specific assumptions on the underlying signals. However, to capture nonlinear time varying, large-scale but also muscle-specific interactions, a more generalised approach is required.

**Approach:** Here we developed a novel framework for muscle synergy extraction that relaxes model assumptions by using a combination of information- and network theory and dimensionality reduction. We first quantify informational dynamics between muscles, time-samples or muscle-time pairings using a novel mutual information formulation. We then model these pairwise interactions as multiplex networks and identify modules representing the network architecture. We employ this modularity criterion as the input parameter for dimensionality reduction, which verifiably extracts the identified modules, and also to characterise salient structures within each module.

**Main results:** This novel framework captures spatial, temporal and spatiotemporal interactions across two benchmark datasets of reaching movements, producing distinct spatial groupings and both tonic and phasic temporal patterns. Readily interpretable muscle synergies spanning multiple spatial and temporal scales were identified, demonstrating significant task dependence, ability to capture trial-to-trial fluctuations and concordance across participants. Furthermore, our framework identifies submodular structures that represent the distributed networks of co-occurring signal interactions across scales.

**Significance:** The capabilities of this framework are illustrated through the concomitant continuity with previous research and novelty of the insights gained. Several previous limitations are circumvented including the extraction of functionally meaningful and multiplexed pairwise muscle couplings under relaxed model assumptions. The extracted synergies provide a holistic view of the movement while important details of task performance are readily interpretable. The identified muscle groupings transcend biomechanical constraints and the temporal patterns reveal characteristics of fundamental motor control mechanisms. We conclude that this framework opens new opportunities for muscle synergy research and can constitute a bridge between existing models and recent network-theoretic endeavours.

## Introduction

The question of whether the control of movement can be characterised by a simplified, low-dimensional strategy is a pertinent one in the motor control literature [1–5]. When one considers the numerous degrees-of-freedom available to the human body at the neural, musculoskeletal and dynamic level for a given movement, the task of selecting an adequate strategy from the set of redundant solutions becomes a computationally intensive operation [3]. Along with this, the capacity for the human body to adapt pre-existing movements and allow for the emergence of novel strategies calls into question, within the context of biomechanical constraints and environmental demands, the nature of the neural constraints on movement [6,7]. It is thought that the central nervous system (CNS) activates movement building-blocks known as motor primitives and through their combination, complex motor patterns can be efficiently performed [3,4]. This strategy allows for the efficient control of groups of neurons, motor-pools and consequently muscles rather than the more computationally intensive, individual control of each degree-of-freedom in the brain. Specifically at the muscle level, evidence for this phenomenon comes most conclusively from animal studies [8,9], but a significant degree of indirect evidence is also accumulating in the human population, for example during development and with training experience [10].

The recent efforts in the motor control literature provide a foundation for furthering our understanding of the mechanisms underlying modularity in human motor control. Among those efforts, different research groups have formulated mathematical definitions to factorize EMG signals using unsupervised machine-learning in the spatial [12], temporal [13] and spatiotemporal domains [14] or their unification through the space-by-time model [15]. These investigations produced novel insights such as the presence of both task-specific and -shared synergies and the complexity of synergies being linearly related to neurological impairment [16,17]. These muscle synergy models are typically implemented using non-negative matrix factorisation (NMF) but tensor decompositions have also recently been utilised with the particular advantage of concurrent extraction of both spatial and temporal synergies and their task-dependent modulations [15,18]. Nonetheless, NMF and its higher-dimensional variants are constrained to extract linear representations of the EMG activity and so may not fully capture the nonlinear characteristics of the musculoskeletal system. Furthermore, the current muscle synergy models don’t facilitate the incorporation of signals with such diverse properties and their extension to include task parameters often violates underlying model assumptions. An example of such an approach is the proposition of ‘functional synergies’ which incorporate task space variables in synergy extraction. Indeed, functional synergies have revealed interesting relationships between muscle activity and biomechanical function [19–21]. However, the muscle- and task spaces are not always expected to share the same mixing coefficients, as task space parameters are not necessarily non-negative and some of the extracted task components may not correspond to EMG data [11,21]. As a consequence of the reliance of these approaches on dimensionality reduction, an emphasis is placed on muscle activations that account for the most variance at the expense of more subtle couplings and task relevancy [22]. Thus, a more generalised and non-parametric formulation of the current muscle synergy models is needed in order to overcome these limitations. Moreover, although current muscle synergy models support recent evidence for multi-functional group membership by individual muscles, [23– 25], the dynamics of these functional groups at the level of pairwise couplings are not well elucidated and may hinder the outputs’ inferentiality. Thus, a methodology that captures pairwise couplings and differential interactions between muscles with respect to a task at multiple spatial and temporal scales may provide more insight into these underlying mechanisms. A generalised approach will also be more amenable to future avenues of research on movement modularity discussed at length by [1], including the falsifiability of muscle synergy patterns by neural signals serving as model constraints. Thus, the development of an appropriate methodological approach that can meet these requirements is incentivised.

In the current study, we sought to develop a novel framework for the characterisation of muscle synergies using a combination of information- and network-theory and dimensionality reduction. Both information- and network theory have proven useful in the analysis of such muscle couplings with novel insights gained in both the temporal and frequency domains and by elucidating the shaping of these couplings by anatomical constraints for example [5,26– 28]. Our proposed framework extends existing muscle synergy models by exploiting the advantageous properties of a novel mutual information formulation for the efficient extraction of salient features across space, time, repetitions and experimental conditions. From the outset, we sought to stringently align the attributes of this framework with the unique characteristics of the synergy concept while positioning it as a useful tool for the progression of the motor control research field. A novel model selection procedure is introduced, where the extracted informational dynamics are modelled as a multiplex network and the predominant clusters are identified. The number of clusters then serves as the empirical input parameter for dimensionality reduction. Following the presentation of this framework, we apply it to two benchmark datasets of point-to-point reaching movements. We identify functionally and physiologically meaningful synergies that demonstrate a high level of consistency in structure, noise correlations and task dependence across participants. A submodular structure representing functional distinct connections is also highlighted across all synergies. This novel architecture of muscle activation signals transcends biomechanical constraints and reveals distributed networks of co-occurring interactions across spatial and temporal scales. Finally, we discuss the continuity of these models with previous research and novel insights gained along with potential directions for future research using this generalised approach. An open-source Matlab GUI (https://github.com/DelisLab/GCMI-synergy-extraction) is available for readers to implement this novel computational framework on their own data as described here.

## Materials and Methods

### Informational dynamics in motor control

A muscle synergy consists of a set of muscles acting as a functional unit during the execution of a coordinated movement and has the identifiable characteristics of a sharing pattern, reciprocal compensation and task dependence [29]. In other words, muscles may share a consistent pattern of activation in space and time, their activations are interdependent and can be adjusted when deviations are experienced, and as elemental variables they can be re-organised for different objective functions. With such complex interdependencies between components, the accurate modelling of these emergent properties during naturalistic behaviour is challenging, requiring a high degree of computational sophistication. This analytical challenge is accompanied by the presence of noisy communication channels that innervate the numerous components of the human nervous system, requiring statistical tools that are robust to noise.

Mutual information (MI) is a statistical measure originally developed to determine the reliability of information transmission across noisy electrical circuits [30]. Recent applications of MI to neural circuitry have been fruitful, with dependence between cortical- and spinal-level activations and motor behaviours elucidated [31–33]. In the following, we briefly present the foundational concepts incorporated in the presented framework that allow for interactions between muscles in space and time to be quantified as informational dynamics in a computationally efficient and noise robust manner.

### Mutual information

The MI between a pair of muscles *M*_*x*_ and *M*_*y*_ (*I*(*M*_*x*_; *M*_*y*_)) (or pair of time-samples *T*_*x*_ and *T*_*y*_ (*I*(*T*_*x*_; *T*_*y*_))) can be thought of as the difference between the entropies of the individual variables (*H*(*M*_*x*_) + *H*(*M*_*y*_)) and their joint entropy (*H*(*M*_*x*_, *M*_*y*_)). The entropy of a random variable is the degree of uncertainty of getting a possible outcome from the given distribution and so MI quantifies the reduction of uncertainty in *M*_*y*_ due to *M*_*x*_.

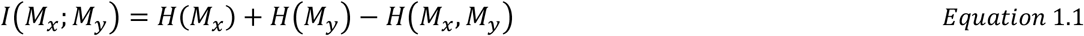

In doing so, MI captures the non-linear dependence between *M*_*x*_ and *M*_*y*_ in the unit known as *bits*. This expression of MI will aid in the communication of the novel MI estimate in the next section. There are a number of equivalent expressions for MI in the literature, another is given in equation 1.2. Here, MI is quantified as the difference between the entropy of *M*_*x*_ and the conditional entropy of *M*_*x*_ given *M*_*y*_, *H*(*M*_*x*_|*M*_*y*_). Using this expression, it can be seen that when *M*_*y*_ completely determines *M*_*x*_ (and vice-versa), the MI is equal to *H*(*M*_*x*_).

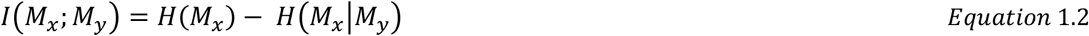

Here we employ a parametric estimator for entropy and MI which enables a reliable estimation of information using limited samples and is computationally efficient [34]. Their formulations are presented in the closed-form in equations 1.3.1 and 1.3.2 respectively. Entropy is expressed as a function of the covariance matrix determinant (|∑|) with dimensionality *k*. 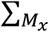 and 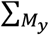 are the covariance matrices of two Gaussian variables (EMG channels, time-sample vectors etc.) and 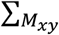 is the covariance matrix for the joint variable contrasted against these individual covariance’s within the determinant. A bias-correction term was applied to these estimations as illustrated in [34].

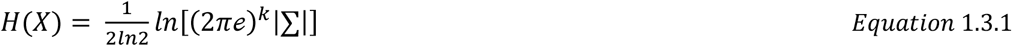

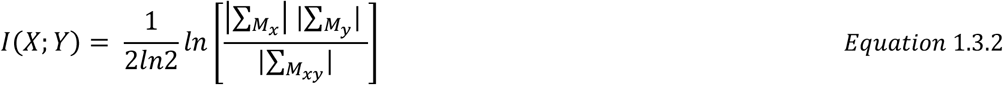

### Gaussian copula mutual information

Similarly to MI, a copula (*c*) is a statistical measure of non-linear dependence between a pair of random variables but in the case of *c*, this estimate has two advantageous properties described in Sklar’s theorem and the invariance theorem [35,36]. Briefly, a given multivariate cumulative distribution function (CDF) can be described by two components, the *c* linking the variables and their marginal CDFs in such a way that when the marginal CDFs are continuous, *c* is unique. *c* is a probability density over the unit square that is formulated through the rank normalization of individual variables and rescaling to a range between 0-1. In linking two random variables, *c* is therefore directly related to MI (Equation 2.1) [34,37].

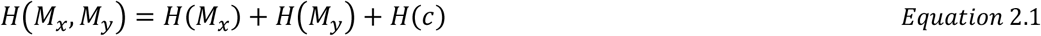

As shown in equation 2.1, the joint entropy of *M*_*x*_ and *M*_*y*_ is equal to their individual entropies and the entropy of *c*. Plugging equation 2.1 into equation 1.1, we see that the marginal entropies will cancel out (Equation 2.2), meaning negative *H*(*c*) is equal to the MI between the *M*_*x*_ and *M*_*y*_ variables referred to hereafter as the Gaussian copula mutual information (GCMI) [34,37].

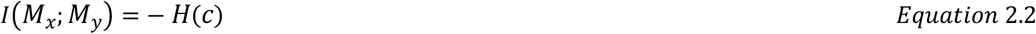

An attractive quality of this formulation is that the empirical copula can maintain this encapsulated relationship between variables following a monotonic transformation of their individual distributions [36]. Thus, it is appropriate to implement the computationally efficient parametric estimation in equation 1.3.1 following a marginal transformation of the individual random variables to a Gaussian distribution. Furthermore, a Gaussian distribution is desirable here as it has the maximum entropy of any distribution and therefore the resulting GCMI estimates serve as a conservative lower-bound on the true MI.

GCMI will form the cornerstone for the muscle synergy framework presented here, representing a robust, nonlinear statistical measure of dependence that can be applied to a broad range of calculations involving uni- and multi-variate samples and in permutation testing for example. An opensource toolbox containing the information-theoretic measures described here was utilised in Matlab software [34].

### Information extraction from EMG activity

To compute couplings between muscles *m* across time *t* = 1, …, *T* (i.e. synergies in the spatial domain), we took a pair of muscle activations *M*_*x*_, *M*_*y*_ for a single trial *s* and determined the GCMI between them *I*_*s*_(*M*_*x*_; *M*_*y*_) (Fig.1 (A)). This procedure was iterated over each unique combination of muscles *k* = 1, …, *K* for all available trials n = 1 … *N*, creating symmetric lower-triangular matrices equal in dimensions to the number of muscles (*M*) in the dataset (Fig.2(A)). These matrices for each trial were vectorised and concatenated (Fig.2 (B)), producing a *N* (No. of trials) x *k*= *M* (*M* − 1)/2 (No. of muscle pairs) matrix 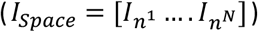 (Fig.2(C)). This matrix served as input into a dimensionality reduction method known as projective non-negative matrix factorisation (PNMF), a variant of the frequently used NMF that has demonstrated a superior capacity for producing sparse representations of high dimensional data and identifying subspace clusters [38,39]. This variant acts as a hybrid of NMF and Principal Component Analysis by using singular value decomposition (SVD) to extract linear, orthogonal features that are positively-constrained. In the current implementation, the SVD was initialised using non-negative Double SVD while the model-rank (i.e. number of extracted components) was determined using a novel procedure (Fig.2(E)) described in detail in the ‘*Model rank selection*’ section that follows [40].

**Fig.1:**
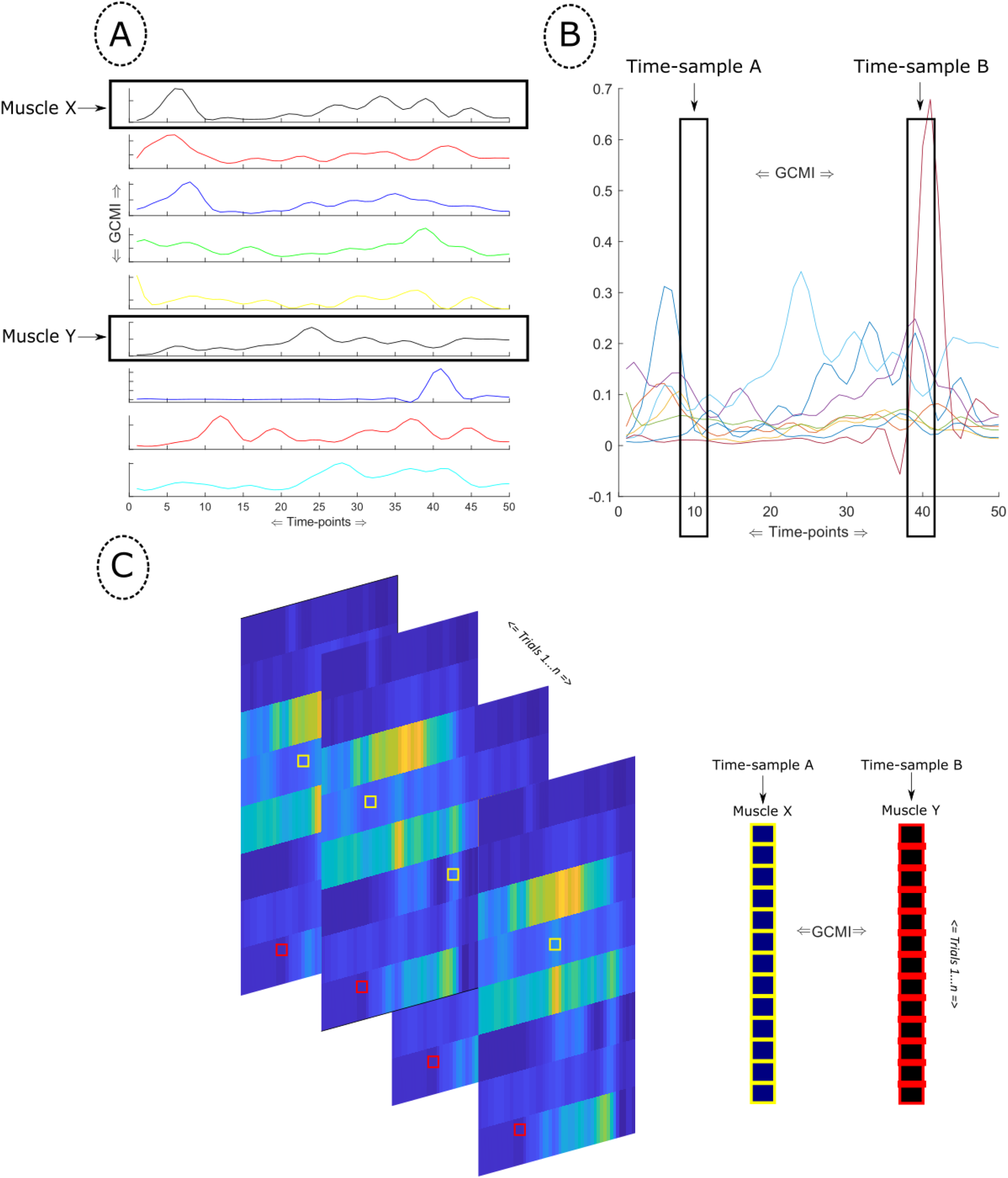
**(A)** An example of spatial information extraction with the GCMI computed from the EMG activity of two muscles across all time-samples in a single trial. **(B)** An example of temporal information extraction where the GCMI is computed between two time-vectors across all nine muscles in a single trial. **(C)** The space-time information extraction is illustrated where time-sample A for a single muscle across all trials is extracted, creating a vector that is used to compute the GCMI against a similar vector for another muscle at time-sample B.

**Fig.2:**
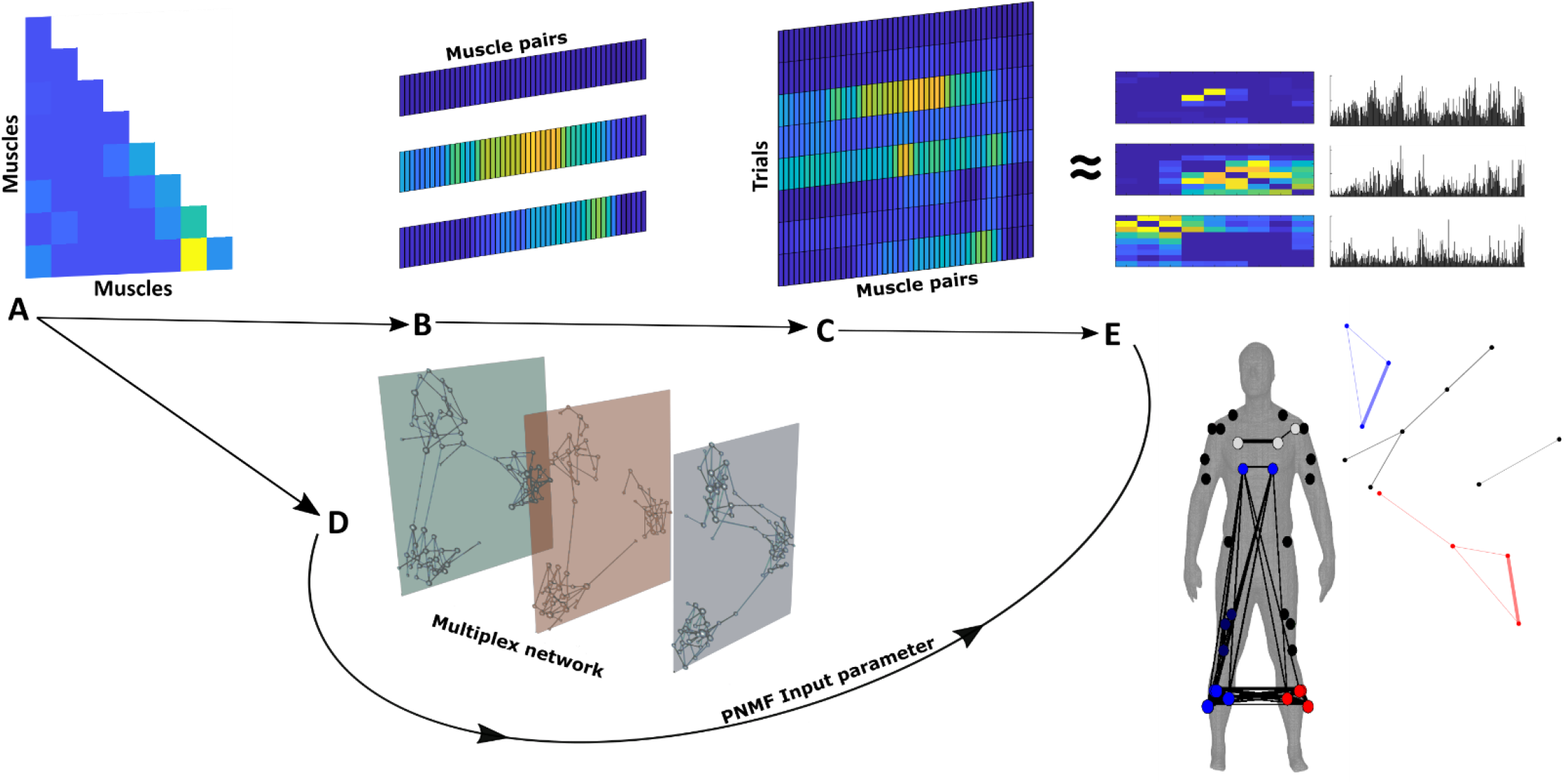
**(A)** The GCMI is computed between each unique pair of muscles (spatial) or time-sample vectors (temporal) or trial-to-trial (space-time) vectors, creating a lower-triangular matrix. **(B)** The lower-triangular matrices are reshaped into single vectors and concatenated, creating the input matrix as shown in **(C). (D)** The matrices depicted in (A) are concatenated into a multiplex network and input into a generalised Louvain algorithm to identify the optimal community structure (See ‘Model rank specification’ section). **(E)** These input matrices shown in (C) are then factorized into lower-dimensional representations of (D) using the optimal community structure as an input parameter to projective non-negative matrix factorisation (PNMF). As an example, we illustrate the spatial synergy output here. The unique structure of the output for each of the synergy models is described in equations 3.1-3.3. A secondary community detection was then conducted to identify submodular structures in the extracted synergies.

In the spatial domain, the factorisation of the input matrix, a matrix consisting of *K* unique muscle pairings across the set of trials *n* = 1 … *N* is described below in vector sum and matrix notation (Equation 3.1). Here the set of synergy weights (*V*) consists of column vectors (*v*) representing clusters of muscle couplings across time in the *jth* module. The set of activation coefficients (*A*) consists of row vectors, 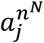, are module- and trial-specific scalar values, inferred to be the neural commands driving the synergy patterns.

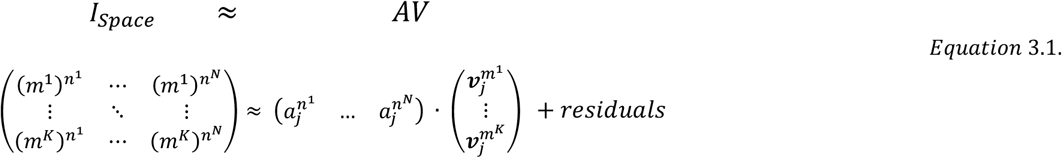

To determine the temporal dependencies across muscles, we computed GCMI between all unique pairs of time-sample vectors of length *m* (L = T(T − 1)/2), creating an *L* x *n* matrix (*I*_*Time*_) (Fig.1(B)). We then implemented the procedure described in Fig.2(A-E), resulting in a factorised output of temporal synergy weights (*W*) and activation coefficients (*A*). The factorised output in the temporal domain is described in equation 3.2 where the set of temporal synergy weights consists of row vectors (***w***) representing clusters of temporal coupling across muscles for the *i*th module.

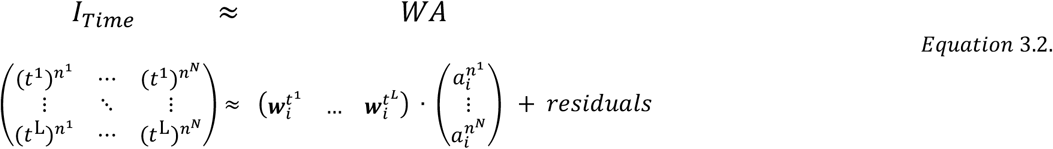

The spatial and temporal models previously described here capture these unique physical dynamics in isolation. To comprehensively capture spatiotemporal dynamics, we developed an extension of the above muscle synergy models and in continuation of previous work [15], spatial and temporal modules were concurrently extracted in a unifying space-time model through the following computation (Fig.1(C)). We extracted the EMG activity of a particular muscle *M*_*x*_ at a particular time-sample *T*_*x*_ across all trials and computed GCMI between this vector and that of a similar vector for muscle *M*_*y*_ at time-sample *T*_*y*_. Once again we iterated this computation over all unique combinations of these vectors and then implemented the synergy extraction procedure described in Fig.2(A-E), i.e. we vectorised the GCMI values computed between these trial-to-trial vectors and concatenated them into a *K* (No. of muscle pairs) x L (No. of time-sample pairs) input matrix (*I*_*Space*−*Time*_). This enabled us to determine trial-to-trial dependencies that are consistent across spatial (*V*) and temporal (*W*) dimensions. The factorised output for the space-time model is described below (Equation 3.3), with ***v*** and ***w*** being the spatial and temporal synergy weights for the *j*th module representing clusters of couplings between muscles in space and time across trials respectively.

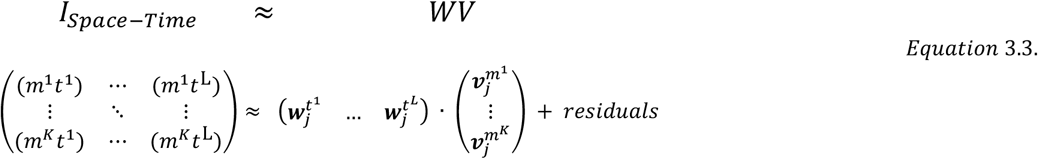

Spurious dependencies typically associated with noise may be found in the above computations and so it was necessary to apply a threshold to identify only the significant associations. A methodology derived from statistical physics, namely a modified percolation analysis [41], was chosen for this framework. This form of percolation analysis was developed to reconcile two opposing characteristics of biological networks, modularity (i.e. the extent of sparsity between groups of densely connected nodes) and ‘small-world’ properties (i.e. the integration of all nodes in a network towards the shortest average path length) [41,42]. The percolation analysis was shown to be effective in identifying functional modular structures while maintaining a sensitivity to the long-range connections that act as the bridges between modules which are often weaker in biological networks [43]. Moreover, this methodology is particularly suited to community detection procedures discussed in the ‘*Model-rank selection’* section that follows [44]. By iteratively removing the weakest dependencies until the *‘giant component’* in the network starts to become affected, the network sparsification threshold can then be defined as this critical stopping point, providing a data-driven method to reduce the effect of noise that can be applied to specific networks.

### Model rank specification

To represent muscle activations adequately, it is necessary to select an optimal number of components to extract. On the other hand, it is also necessary to maintain a model rank that produces physiologically and functionally meaningful synergies while also identifying a low-dimensional space, a challenging task within the literature [45]. To accommodate the unique characteristics of the GCMI matrices generated here, we developed a novel model rank selection criterion by defining a muscle synergy, a group of muscles acting cohesively as a functional unit, in network-theoretic terms. More specifically for the identification of these functional units in our data, we defined a synergy as a densely connected set of nodes otherwise known as a community [46]. Hereafter, the term node refers to either a muscle, time-sample or muscle - time-sample pairing depending on the model in question. Prior to dimensionality reduction using PNMF, the row for each trial of an input matrix was converted to an adjacency matrix and became a layer in a multiplex network with all other trials. In the case of space-time computations, the temporal pattern across each spatial vector constituted the multiplex network. We implemented the percolation analysis described previously on each individual layer of the supra-adjacency matrix before the following computation.

Modularity in a monoplex network can be determined by the difference between the sum of all vertices that fall between all possible pairs of vertices in the network and that which would be expected by random chance [42]. This quantity is known as the Q-statistic which acts as an objective function to be optimised in community detection algorithms, ranging from 0-1 with 1 indicating maximal modularity [42,47]. The multiplex modularity, a generalisation of the Q-statistic to multi-layered networks, was implemented here using the generalized Louvain algorithm where the community structure of intra-layer edges between all nodes and all inter-layer edges between nodes representing the same muscle/time-sample could be considered [46,48]. The resolution parameter γ was set to the default value of 1 for classical modularity [46]. The number of communities identified served as the input parameter for PNMF (Fig.3(E)). To verify the communities identified a priori were in fact extracted using PNMF, the maximum value in each row across synergy weight vectors (***w***) served as an indicator for hard-cluster assignment [38]. By visually inspecting the cluster assignments produced by PNMF and those found through community detection, the output was found consistent (see supplementary materials (Fig.1.) for an example of this procedure).

**Fig.3.**
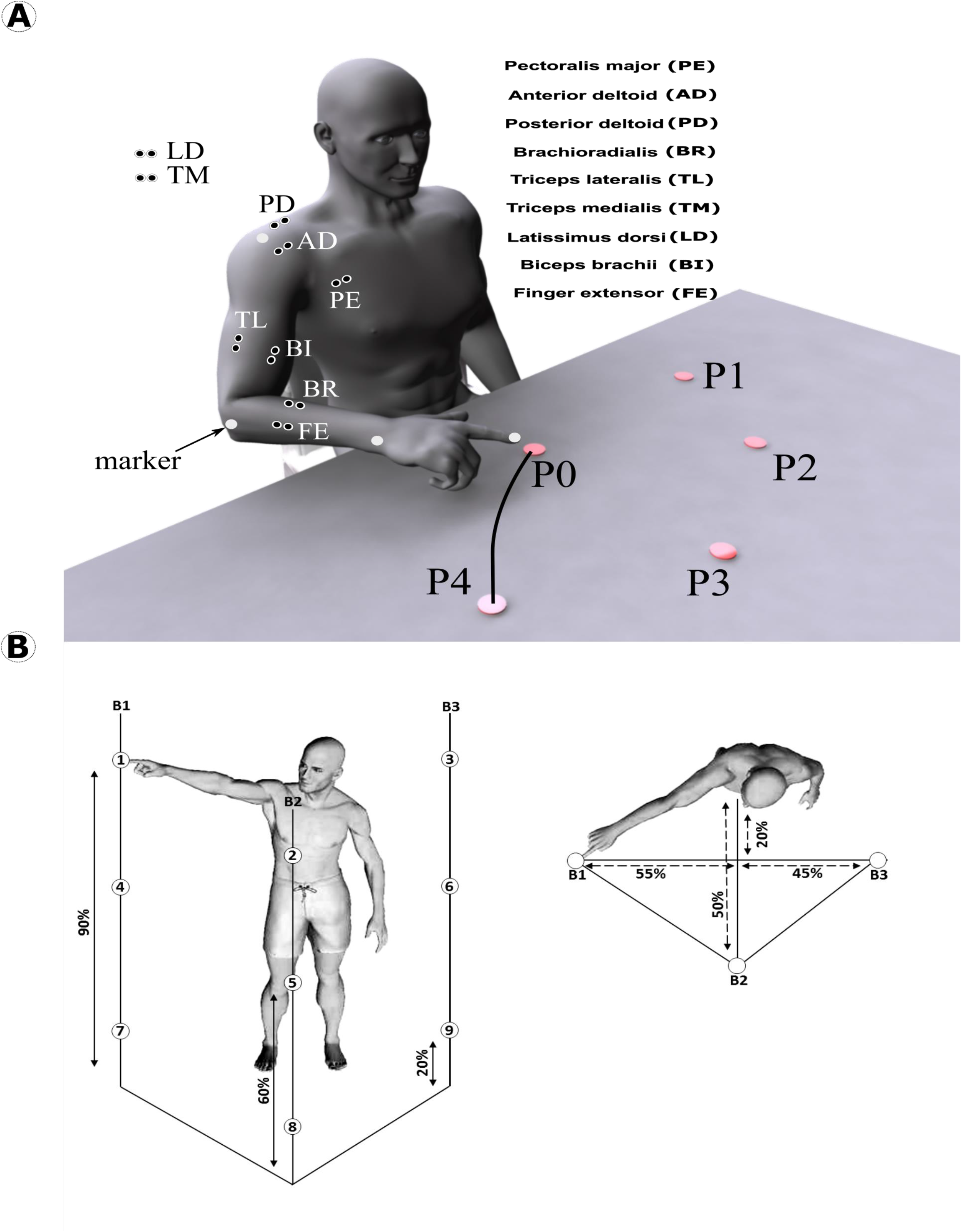
(A): An illustration of the experimental setup for Dataset 1 [14]. 40 repetitions for each target (P1-P4) were conducted while the EMG activity from the following muscles was recorded: LD: Latissimus dorsi; AD: Anterior deltoid; PD: Posterior deltoid; PE: Pectoralis; TL: Triceps lateral; BI: Biceps brachii (long head); TM: Triceps medial; BR: Brachioradialis; FE: Finger extensors. **(B):** An illustration of the experimental setup for Dataset 2 [57]. Participants were asked to perform whole-body pointing movements between pairs of targets. Included were 9 different targets at 3 different heights, creating 72 possible movement tasks for analysis.

### Trial-to-trial network configuration and task dependence of extracted muscle synergies

Following the extraction of muscle synergies using the GCMI framework, we then investigated whether the identified sharing patterns also aligned with the other attributes of the synergy concept, i.e. reciprocal compensation/flexibility and task dependence [29]. As mentioned in the previous section, biological networks are generally characterised by two competing properties, fractal modularity and small-worldedness [41,43]. The scale invariance of network architectures allows for greater robustness to injury and adaptability while the efficient transfer of information is crucial for within- and across-network communication and cohesion [49,50]. It is therefore unsurprising to find modularity has been related to the motor learning process and small-worldedness to important integrative mechanisms such as working memory performance [51,52]. In the context of the synergies extracted here and their reorganisation, fluctuations in these network properties from trial-to-trial may provide interesting insight into the neural control of movement. This is supported by experimental findings where increased integration in the muscle space and the reorganisation of modular structure from trial-to-trial were observed with additional motor noise and adaptations to task constraints respectively [53,54].

Thus, we sought to determine if the GCMI framework could capture the dynamic balance between modularity and small-worldedness from trial-to-trial in the extracted activation coefficients. We firstly determined the global efficiency (GE) and Q-statistic for modularity (Q) for individual layers of the thresholded multiplex network depicted in Fig.3(D). GE quantifies the small-worldedness of a network through the ratio of present to possible edges [55]. Q was determined for individual network layers using the conventional Louvain algorithm [47,55]. Analogous to previous work [56], this created two vectors in the spatial and temporal domains that were demeaned with respect to the task along with their corresponding activation coefficients. The correlations between single trial synergy activations and the network property measures defined above (GE, Q) for a fixed task (noise correlations) were then computed using Pearson’s correlation [56]. The strength and direction of these correlations therefore provide insight into the sensitivity of the extracted synergies to this network level trade-off. To further exemplify the fractal modularity of the synergies produced by this framework, a secondary community detection procedure was carried out on the synergy weights using the traditional Louvain algorithm [47,55]. The cluster assignment of these submodules were then represented on a human body model by the colour of nodes on the network [57].

To determine if the information encoded in the extracted muscle synergies can discriminate between the different experimental conditions/tasks, we computed the GCMI between individual spatial or temporal activation coefficient vectors and the reaching task variable. To assess whether the task-discriminative information presented in each component was significant, activation coefficients were also randomly shuffled and the GCMI with respect to the task was calculated. This procedure was repeated 100 times and the 95^th^ percentile of these collective values acted as an empirical threshold for significance. The specific task attributes for each dataset analysed in this study are detailed in the ‘*Experimental design and setup’* section.

### Structural similarity of the muscle synergies across participants

It is crucial to the utility of this framework that the muscle synergies extracted are consistent across a set of participants who have conducted the same motor tasks. The same extraction procedures were therefore conducted across the remaining participants in each dataset and the following analysis was conducted to determine the structural similarity. Muscle synergies were first visually inspected and compared across participants and those with a similar functional interpretation were paired together. Following their vectorisation, the Pearson’s correlation was then found between each pairing. This produced a set of coefficients that were transformed into Fisher’s Z values, the average and standard deviation of which was taken and reverted to correlation coefficients. This value served as a global index of similarity across participants. To extract the average muscle synergy from each model, the synergies across participants in each dataset were vectorized and the mean network was then found. To eliminate individual differences from the average synergies, the percolation analysis sparsification procedure described above was conducted.

### Experimental design and data collection

#### Dataset 1

A dataset of EMG activity recordings was generated across seven healthy, right-handed adults (Age: 27±2 years, Height: 1.77±0.03m) who provided informed consent to participate in the experiment that conformed to the declaration of Helsinki (approved by the local ethical committee ASL-3 (“Azienda Sanitaria Locale,” local health unit), Genoa). Participants were instructed to perform center-out (forward, denoted by *fwd*) and out-center (backward, denoted by *bwd*) one-shot point-to-point movements between a central location (P0) and 4 peripheral locations (P1-P2-P3-P4) evenly spaced along a circle of radius 40 cm at a fast pace (370-560 m/s duration on average across participants). 40 repetitions per target/speed were captured per participant using the dominant hand. Movement onset and offset phases were defined at the points in which the velocity profile (captured via the kinematic data of a passive marker (Vicon, (Oxford, UK)) placed on the finger-tip that was numerically differentiated) fell above/below 5% of its maximum. The targets to which the participants reached were circles of radius 2 cm which they had to touch. No restraints or supports were provided while the order of the tasks were randomized.

Simultaneously, the activity of nine upper-body and arm muscles was recorded as presented in Fig.3(A) (Aurion (Milan, Italy)). The EMG setup was implemented in line with SENIAM guidelines. EMG signals were digitized, amplified (20-Hz high-pass and 450-Hz low-pass filters), and sampled at 1,000 Hz (synchronized with kinematic sampling). Subsequently, to extract the signal envelopes, the EMG signals were processed offline using a standard approach [14]: the EMGs for each sample were digitally full-wave rectified, low-pass filtered (Butterworth filter; 20-Hz cut-off; zero-phase distortion), normalized to 1,000 time-samples and then the signals were integrated over 20 time-step intervals yielding a waveform of 50 time-steps. Finally, the EMG signal of each muscle was normalized in amplitude by dividing each single-trial muscle signal by its maximal value attained throughout the experiment. For dataset 1 participants, the task dependence of individual spatial and temporal ***a*** was determined against discrete variables representing all point-to-point movements (P1-P8), all center-out (forward) movements (P1-P4), all out-center (backward) movements (P5-P8) and the movement direction (fwd vs. bwd).

#### Dataset 2

The EMG activity of 30 muscles from four healthy, right-handed adults (Age = 25 ± 3 years, height = 1.72 ± 0.08 meters, weight = 70 ± 7 kg) was recorded (Aurion system, (Milan, Italy), SENIAM guidelines) while participants performed whole-body point-to-point movements in various directions and at differing heights. The muscles recorded included: tibialis anterior, soleus, peroneus, gastrocnemius, vastus lateralis, rectus femoris, biceps femoris, gluteus maximus, erector spinae, pectoralis major, trapezius, anterior deltoid, posterior deltoid, biceps and triceps brachii on both sides. The experimental setup (Fig.3(B)) was approved by the Dijon Regional Ethics Committee and conformed to the Declaration of Helsinki. Written informed consent was obtained by the participants following guidelines set by the Université de Bourgogne. This setup is described in detail elsewhere and will therefore be outlined briefly here [58]. Participants were asked to perform whole-body pointing movements between pairs of targets (9 targets at three different heights = 72 different movement tasks). Over the course of two separate sessions, participants performed 15 repetitions of each movement (30 repetitions in total), resulting in 72 × 30 = 2160 trials per participant. The processing of EMG recordings followed the same standardized protocol conducted on Dataset 1 while onset and end times for individual trials were determined in a similar manner using the kinematics of the pointing index finger at a frequency of 100 Hz [14,59]. The left arm of participants was unconstrained and at rest throughout each trial while the order of movements was randomised. The resulting data was pooled, creating (50 time-sample x 2160 trials) x 30 muscle matrices. For dataset 2 participants, the task dependence of individual spatial ***a*** was determined against discrete variables representing the spatial movement features, i.e. start- and end-point bar and height (see fig.3(B)), up-down and left-right reaching directions. For individual temporal ***a***, the task dependence was determined with respect to temporal movement features representing the evolution of movement in time determined with respect to temporal movement features representing the evolution of movement in time, i.e. start-point, reaching direction and end-point target.

## Results

### Muscle synergies in point-to-point reaching movements

#### Spatial synergy model

To identify spatial synergies, i.e. muscle couplings across time within each trial (see Fig.1(A)), we applied the proposed framework to dataset 1 consisting of arm-reaching movements in 8 different directions. In brief, we first computed GCMI between all pairs of muscles for each trial. We then input the single-trial GCMI values of all muscle pairs to the PNMF algorithm to identify couplings between muscles, i.e. muscle synergies, which are consistent across trials (see Fig.2 for an illustration of the full methodology).

Taking a representative example participant, three distinct spatial synergies were found using the community detection procedure (see Materials and Methods for details) with a Q-statistic value of 0.998 (Fig.4(A)). A high co-activation between the medial and lateral triceps was found in S1 along with a lower co-activation of the anterior deltoid. S2 represented a more global activation across the upper-arm and shoulder muscles while the finger extensors and brachioradialis remained inactive. S3 contained couplings between the finger extensors, brachioradialis and biceps brachii. Taking these together, one can interpret their functional grouping as the following: S1 represents elbow extension, S2 proximal stabilisation/shoulder flexion, and S3 is forearm flexion/finger control.

To assess the task-relevance and organisation of the identified synergies, we computed the task information conveyed by the synergy recruitment on each trial (Fig.4(B)) and the noise correlation between synergy recruitment and essential network properties derived prior to dimensionality reduction (Fig.4(C)). Forward vs. backward movements were mainly discriminated by S2 activations (0.26 bits) which implement shoulder flexion. S1 (elbow extension) contained the most information for movement direction across all movements, i.e. P1-P8 (0.51 bits), but also when considering only forward (P1-P4, 0.45 bits) or only backward (P5-P8, 0.52 bits) movements. S3 activations (predominantly of forearm muscles) did not contribute significantly to discrimination of forward movements (P1-P4, 0.16 bits) or forward vs backward direction (0.033 bits) but carried significant information for backward movements (P5-P8, 0.34 bits) which contributed to the overall discriminatory information (P1-P8, 0.29 bits). We also found significant correlations between all three synergy activation coefficients and network-theoretic measures (p<0.05) but in opposing directions with positive associations for integration (GE) and negative associations for segregation (Q). Both S2 and S3 shared the highest association with GE (R= 0.72), an interesting finding considering these synergies represent the distinct functional grouping of upper- and lower-arm muscles respectively. The activation in S1 reduced the least with positive changes in modularity (R= -0.11), likely due to the specificity of this synergy to the medial- and lateral-triceps.

**Fig.4:**
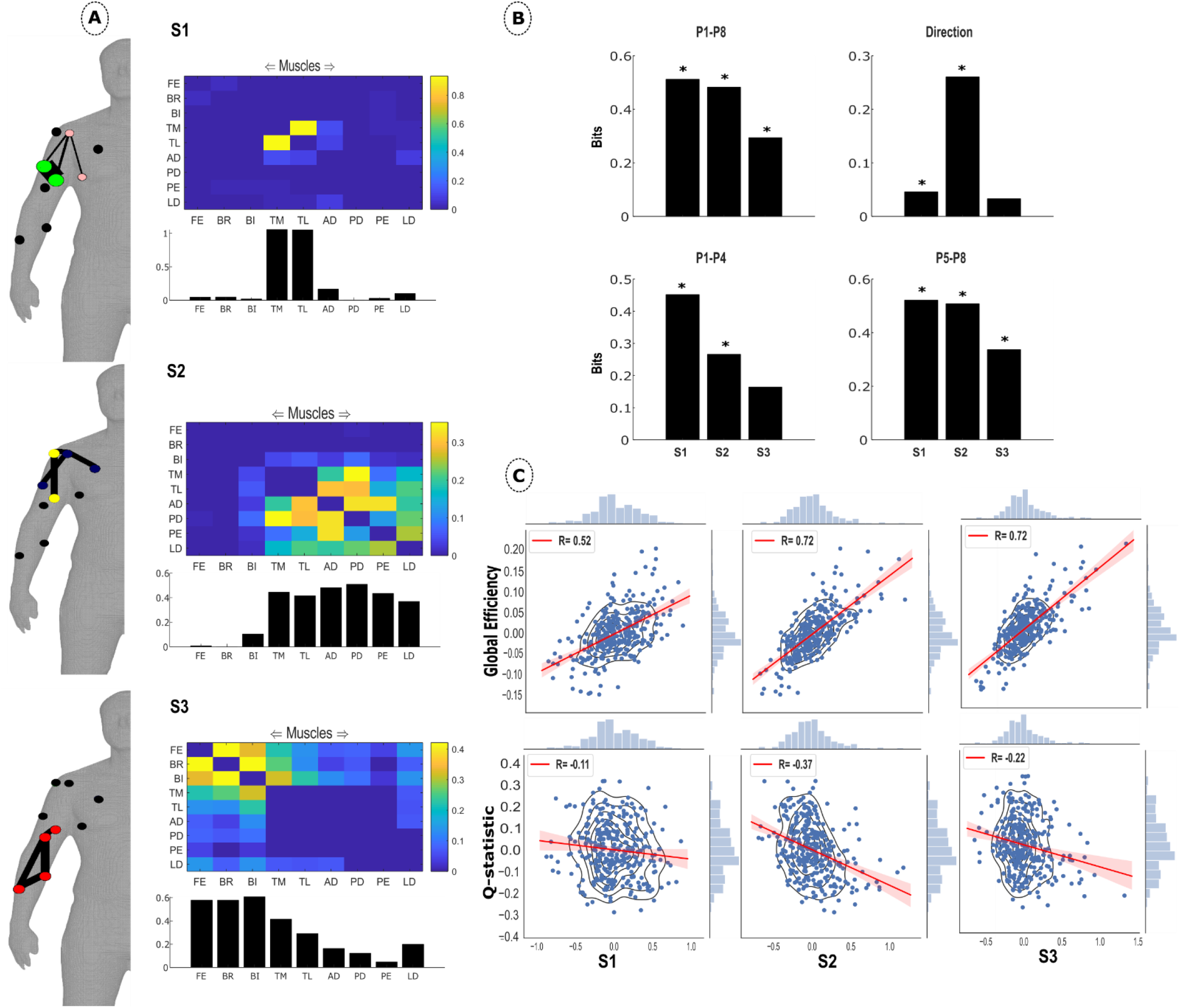
**(A)** The spatial synergies extracted from an example participant in Dataset 1. The community detection criteria found three distinct communities that were extracted using PNMF. The bar graph below represents the average values in each column of the adjacency matrix. The minimally connected human body model illustrates the values in the adjacency matrix with the width of the edges and colour and size of the nodes providing insight into connection strengths, submodular structure and involvement respectively [57]. Submodular structure was identified using the conventional Louvain algorithm on the synergy matrices [47]. Unconnected nodes are in black. **(B)** The encoded information of the three spatial synergies identified in the example participant in dataset 1 for four task attributes: P1-P8, P1-P4, P5-P8 and Direction. Significant information (p<0.05) is indicated with *.**(C)** The correlation between trial-to-trial fluctuations in Global Efficiency/Q-statistic for modularity and noise in the spatial activation coefficients. The bars along the axis of each plot are marginal histograms of the x- and y-variables.

A secondary community structure analysis of the extracted synergies revealed two submodules in S1 and S2 and just one submodule consisting of the lower-arm musculature in S3. In S1, these submodules consisted of the medial- and lateral triceps (green) and anterior deltoid and latissimus dorsi (pink). The medial-triceps and posterior deltoid (yellow) and lateral-triceps, pectoralis major and anterior deltoid (navy) comprised the S2 submodules. These submodules represent the distinct (but often overlapping) co-occurring muscle couplings that contribute consistently across tasks/trials to single-joint or cross-joint actuations.

#### Temporal synergy model

To characterise the temporal structure of muscle co-activations, we also applied the GCMI framework in the temporal domain, returning three temporal synergies (Fig.5(A)) for the same example participant that captured temporal associations across muscles. This was produced by computing GCMI between pairs of time-vectors (vectors of activations across all muscles at different time-samples, see Fig.1(B)) within each trial, modelling the output as a multiplex network to identify the optimal community structure and finally extracting these communities using PNMF.

T1 presented dependencies between time-samples for the initial phase of the reaching movement. These dependencies are accompanied by a high dependency between the final (49^th^-50^th^) time-samples that is connected to this initial burst through moderate dependencies among adjacent time samples along the diagonal. T2 illustrates a prominent clustering of dependency from time-samples 15-25 and continuing with strong connection strengths within this time-sample range for the remainder of the movement. T3 presented high dependencies with all time-samples from approximately half-way into the movement until the endpoint.

**Fig.5:**
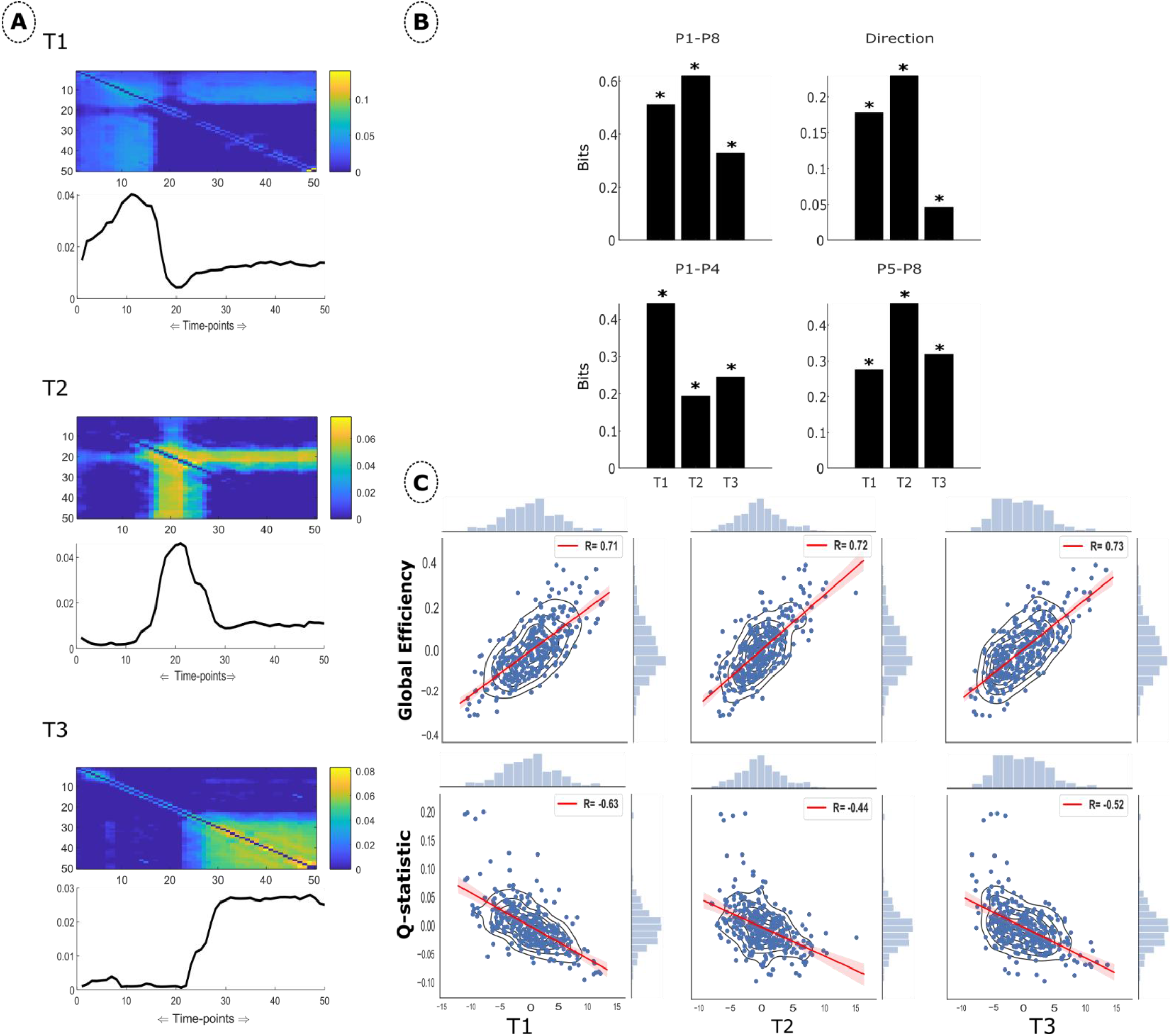
**(A)** The temporal synergies from the example participant in dataset 1. The community detection criteria found three distinct communities that were extracted using PNMF. The line plot below represents the average values within each column of the adjacency matrix. **(B)** The encoded information of the three temporal synergies identified in the example participant in dataset 1 for four task attributes: P1-P8, P1-P4, P5-P8 and Direction. Significant information (p<0.05) is indicated with *. **(C)** The correlation between trial-to-trial fluctuations in Global Efficiency/Q-statistic for modularity and noise in the temporal activation coefficients. The bars along the axis of each plot are marginal histograms of the x- and y-variables.

Taken together, a single tonic synergy (T1) was accompanied by a bi-phasic pattern of activation (T2 (acceleration) and T3 (deceleration)) that explained point-to-point reaching movement in this example participant. This observation is supported by an analysis we conducted on the task-dependency of the underlying activation coefficients (Fig.5(B)). Consistent with the differing functional roles of these temporal synergies, differences between forward movements (P1-P4 differing in the respective end-points) were predominantly captured by T1 (peaking at the final time-samples) whereas differences in backward movements (P5-P8 differing in the respective starting points) were predominantly captured by T2 (activating earlier in the movement) which contained the most information overall (P1-P8, 0.62 bits and direction, 0.23 bits). The noise correlations between all synergy activations and both GE and Q were significant (p<0.05) and in opposing directions, with all synergy activations positively related to GE and negatively associated with Q (Fig.5(C)). The tonic synergy (T1) was most sensitive to changes in modularity from trial-to-trial (R= -0.63), while T3 was highest in its association with GE (R= 0.73). The sensitivity to integration in the latter case is likely reflective of the differing muscular involvements required at each reaching end-point position.

### Space-Time synergy model

As an extension of the spatial and temporal synergy models, we concurrently extracted these distinct modules in a unifying space-time model, capturing distinct couplings in trial-to-trial dependencies that are consistent across spatial and temporal components. To do this, we first computed GCMI across all muscle pairs and time-samples and then decomposed the identified dependencies into components with distinct spatial and temporal signatures. For the same example participant in dataset 1, we determined an optimal model-rank of three using the community detection procedure with a Q-statistic of 0.97. Fig.6 below illustrates this output with the spatial and temporal synergies corresponding on a 1:1 basis across each row. In other words, the space-time model identified three synergies and characterised their spatial and temporal structure across all trials. The spatial synergies in ST2 and ST3 are relatively robust with those derived from the spatial synergy model previously (Fig.4), demonstrating the consistent coupling of the medial and lateral triceps and within the upper- and lower-arm musculature respectively. Their corresponding temporal synergies represent muscle co-activation throughout the movement and an early-and-late phasic activation that is dynamically involved in the flexion/extension of reaching movements respectively. ST1 presents a strong dependency between the anterior deltoid and latissimus dorsi that is active at the beginning and end of the movement and relatively quiet in the mid-range. This synergy is likely reflective of the alternate roles the anterior deltoid and latissimus dorsi carry out in stabilising the shoulder joint during forward and backwards reaching movements.

**Fig.6:**
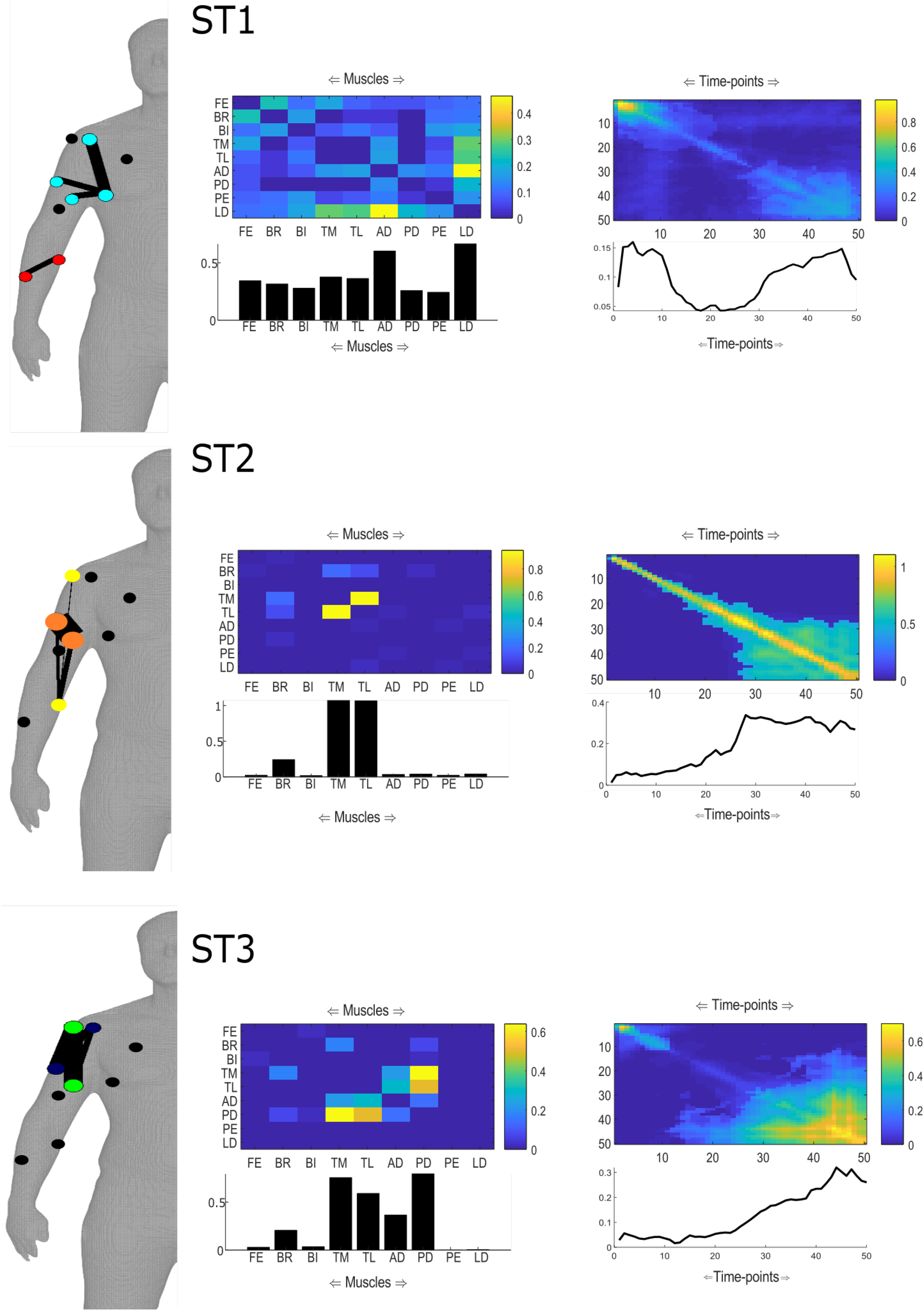
The Space-Time synergies extracted from the example participant in dataset 1. Three communities were identified a priori and then extracted using PNMF. Spatial and temporal synergies correspond on a 1:1 basis here as presented across rows. The bar/line plots below represents the average values within each column of the corresponding adjacency matrix. The connection strengths, submodular structure and involvement of nodes are indicated by the minimally connected human body model via the edge widths, node colour and size respectively [57]. Submodular structure was identified using the conventional Louvain algorithm on the synergy matrices [47]. Unconnected nodes are in black.

A secondary community detection procedure revealed two submodules in each of the spatial synergies that are represented by the colours of individual nodes in the human body model networks. For ST1, the submodules included the anterior deltoid, latissimus dorsi and both the medial- and lateral triceps (cyan) and the finger extensors and brachioradialis of the lower-arm (red). In ST2, the medial- and lateral triceps (orange) and a long range connection between the brachioradialis and posterior deltoid (yellow). Finally, ST3 consisted of the medial-triceps and posterior deltoid (blue), and lateral-triceps and anterior deltoid (green). Both ST1 and ST3 submodules were anatomically compartmentalised and non-overlapping, indicating localised functionalities that were co-occurring and reproducible across trials such as joint stiffness and stabilisation. ST2 submodules represented a more global interaction between overlapping submodules across the arm, coinciding with its corresponding temporal synergy representing a co-activation across the entire movement.

### Consistency across participants (Dataset 1)

We then sought to determine the similarity of the spatial, temporal and space-time synergies extracted using the GCMI framework across participants. This was conducted by computing a similarity index between pairs of functionally similar synergy weights derived at a representative model rank for the dataset and the average ± standard deviation was found (see Materials and Methods section). Within Dataset 1, we identified 2.4 spatial communities on average (range= 2-3) with a mean Q-statistic of 0.997 and threshold value of 0.13 bits (supplementary materials (Fig.2)). A satisfactory level of consistency was found (R=0.78±0.37). S1 here represents elbow extension primarily with some moderate dependencies across the other muscles. S2 involves a more global activation but particularly between the medial and lateral triceps and between the anterior deltoid and pectoralis. S1 was typically higher in its task information across participants for direction (0.11 bits) and P5-P8 (0.23 bits) while S2 was highest for P1-P8 (0.33 bits) and P1-P4 (0.24 bits). The contrasting relationship between the synergy activations and GE and Q was replicated across participants with S1 presenting the strongest noise correlation for both network properties (supplementary materials (Fig.2(C)).

We identified an average of 3.4 communities (range= 3-4) with an accompanying maximal modularity of 0.93 and threshold value of 0.46 bits on average in the temporal domain across participants in dataset 1. The similarity index revealed a high level of correlation (R=0.83±0.45) although with significant variability. Supplementary materials (Fig.3(A)) illustrates the three representative synergies taken from this procedure along with their mean task-encoded information. T1 presented a low-to-moderate level burst of dependency at the initial phase of movement, but a high dependency in the last two time-sample pairings. T2 had a more idiosyncratic pattern of dependencies across the movement with several bursts along the diagonal. T3 demonstrated a step in activation from the 30^th^ time-sample approximately with dependencies shared across time-samples for the remainder of the movement. In terms of task dependence (supplementary materials (Fig.3(B)), T1 was predominant for P1-P8 (0.31 bits) and P1-P4 (0.22 bits) and also highest for P5-P8 (0.23 bits). T2 contained the highest average task information for forward vs. backward directions (0.11 bits). T1 demonstrated the strongest correlation for both GE (R= 0.86) and Q (R= -0.57) while T3 was weakest in its noise correlation with Q (R= -0.16) (supplementary materials (Fig.3(C)).

For the space-time model, we determined that a model rank of three was representative of dataset 1 participants with a correlation of 0.69±0.33 found (supplementary materials (Fig.4)). The Q-statistic was consistently high across participants, ranging from 0.89-0.99 while the mean threshold value was 0.048 bits. All temporal synergies presented a high dependency between adjacent time-samples and along the diagonal, indicating that co-activations during reaching movements in various directions and points were most consistent across participants when considering trial-to-trial dependencies. The spatial synergies were consistent with those reported already here with ST2 capturing the characteristic dependency between the medial and lateral triceps and the ST3 spatial synergy the dynamic involvement of the upper-arm musculature during reaching. ST1 involved a combination of the anterior deltoid and pectoralis or latissimus dorsi.

### A generalisation to whole-body point-to-point reaching movements

We then sought to generalise the results presented above to a more complex dataset consisting of EMG activity from 30 muscles during whole-body point-to-point reaching movements at various heights and in various directions (82 distinct movements in total, each repeated 30 times) among 5 participants. This new high-dimensional dataset serves to demonstrate the applicability of the proposed framework to characterise the structure of large-scale EMG recordings during 3-dimensional unconstrained movements.

#### Spatial synergy model

Taking an example participant from dataset 2, we identified three spatial synergies with a maximal modularity of 0.994 (Fig.7(A)). S1 and S2 appear to represent the postural stabilisation activity related to normal stance and during point-to-point reaching respectively. This is indicated in S1 by the greater dependencies found in the lower-limbs and in the lower-limbs and left upper-body in the S2. S3 then completes this picture of whole-body reaching with significant dependencies clustered in the right upper-body. The dependency of these synergies across a number of task attributes including end-point bar and height, start-point bar and height along with the Up-down and left-right directions are also illustrated (Fig.7(B)), and support these functional interpretations. S1 (activating mainly the lower body) carried significant information about the horizontal dimension of movement (start and endpoint bar and left-right displacements) suggesting that its functional role was to drive body rotations. S2 (activating the left upper-body together with parts of the lower body) contained a higher level of task information, mostly for the vertical movement dimension (end-point height [0.242 bits] and the Up-down direction [0.208 bits]) suggesting that its functional role was to support vertical body displacements. Although task information was relatively high for S3 (right arm raising), just one task attribute was found to be significant (end-point height, 0.53 bits), suggesting that this synergy was relevant but highly variable in the information it contained. This could potentially be attributed to the highly variable demands required during upwards vs. downwards reaching movements for example where passive mechanics can be exploited in the latter but not the former. All synergy activations were significant in their trial-to-trial correlations with integrative and segregative network properties (p<0.05), and presented contrasting directions in their association, exemplifying the trade-off between these properties. S2 demonstrated the greatest sensitivity to changes in GE (R= 0.62)and Q (R= -0.41) in its underlying activations. Coinciding with the highly variable task information, S3 presented the least sensitivity to changes in GE (R= 0.29) and Q (R= -0.07).

**Fig.7:**
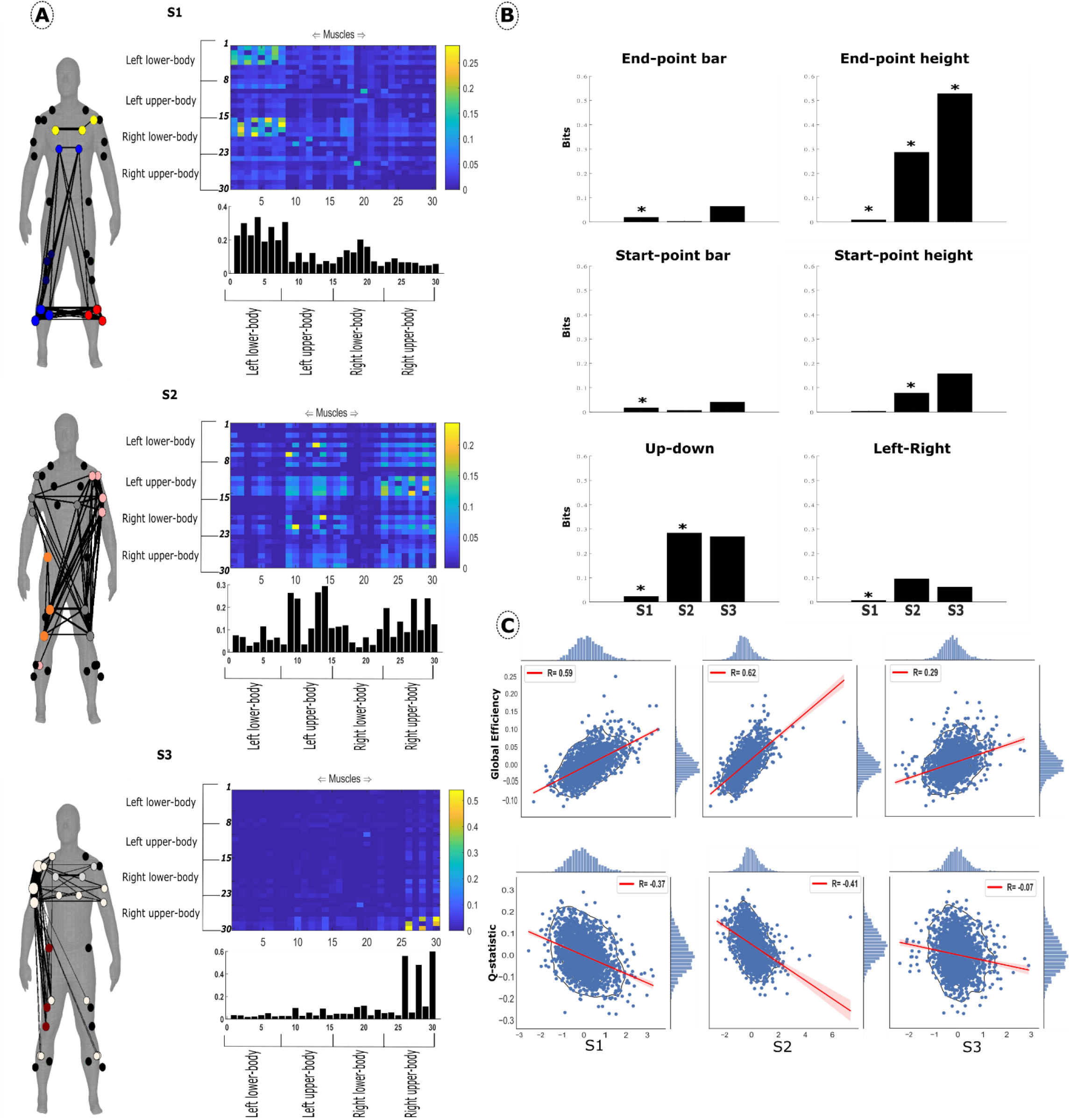
**(A)** The spatial synergies extracted from the example participant in dataset 2. The adjacency matrices are organised so that rows 1-8: left lower-limb, rows 9-15: left upper-body, rows 16-23: right lower-limbs and rows 24-30: right upper-body. The width of the edges on the minimally connected human body model and the size and colour of the nodes indicate the connection strength, node involvement and submodular structure respectively [57]. Submodular structure was identified using the conventional Louvain algorithm on the synergy matrices [47]. Unconnected nodes are in black. **(B)** The information encoded for six task attributes is presented. * indicates significance at p<0.05.**(C)** The noise correlation between spatial synergy activations and trial-to-trial network properties Global Efficiency/Q-statistic for modularity. The bars along the axis of each plot are marginal histograms of the x- and y-variables.

The submodular structure of these synergies, represented on the human body model by the colour of the nodes shows four submodules for S1 (both erector spinae and the right tibialis musculature (blue), right femoral musculature (navy), left tibialis musculature (red) and both pectoralis major and the left anterior deltoid (yellow)) three submodules for S2 (left arm musculature and right tibialis anterior (pink), right arm and left femoral musculature (grey) and right femoral musculature (orange)) and two submodules for S3 (upper-body musculature (white) and three right femoral muscles (maroon)). The community assignment of these distributed networks provides further evidence as to the underlying function of specific muscle couplings. For example, an interesting long-range connection is found between the right-side lower-limb musculature and erector spinae that belong to the same submodule (S1). This coincides with an overlapping but distinct community among the left-side lower-limb musculature, indicating that the non-reaching side contributes differently to whole-body stability than the reaching side. The predominant submodule in S3 is specific to the reaching arm where a large cluster of strongly weighted connections are present. Long-range connections to distal body parts are also evident in S3, indicating that the whole-body is coupled to the specific activations of the reaching arm.

#### Temporal synergy model

To test how consistent the temporal synergies are when considering more complex motor tasks, we applied the temporal synergy model to the example participant in dataset 2, revealing three temporal synergies (Fig. 8(A)). T1 here contained moderate proximal dependencies along the diagonal and a gradual increase to a new activation level near movement termination, characteristic of a tonic activation. The initial phasic activation of T2 here is more drawn out along the diagonal and less dependent on time-samples later in the movement. This is likely due to the different heights and directions in point-to-point reaching introduced in this experimental setup, requiring more variable onset-offset timing of phasic muscle activations. T3 consists of a late burst from time-sample 30-40 approximately. We found that all of these temporal synergies were significant for the three task attributes analysed here including start- and end-point and direction (Fig.8(B)). T3 contained the most information regarding the direction of reaching (0.13 bits). All 3 synergies contained similar amount of information about end-point position (0.11, 0.1 and 0.11 bits respectively) while T2 was predominant for start-point position (0.1 bits). The dynamic balance between modularity and small-worldedness was once again captured in the extracted activations (Fig.8(C)), with opposing directions of correlation in the trial-to-trial fluctuations for GE and Q that were all significant (p<0.05). T3 activations were most strongly related to integrative changes (R=0.75), once again likely reflecting the differing muscular involvement required at end-point position while T2 demonstrated the strongest association with trial-to-trial changes in modularity (R= -0.83).

**Fig.8:**
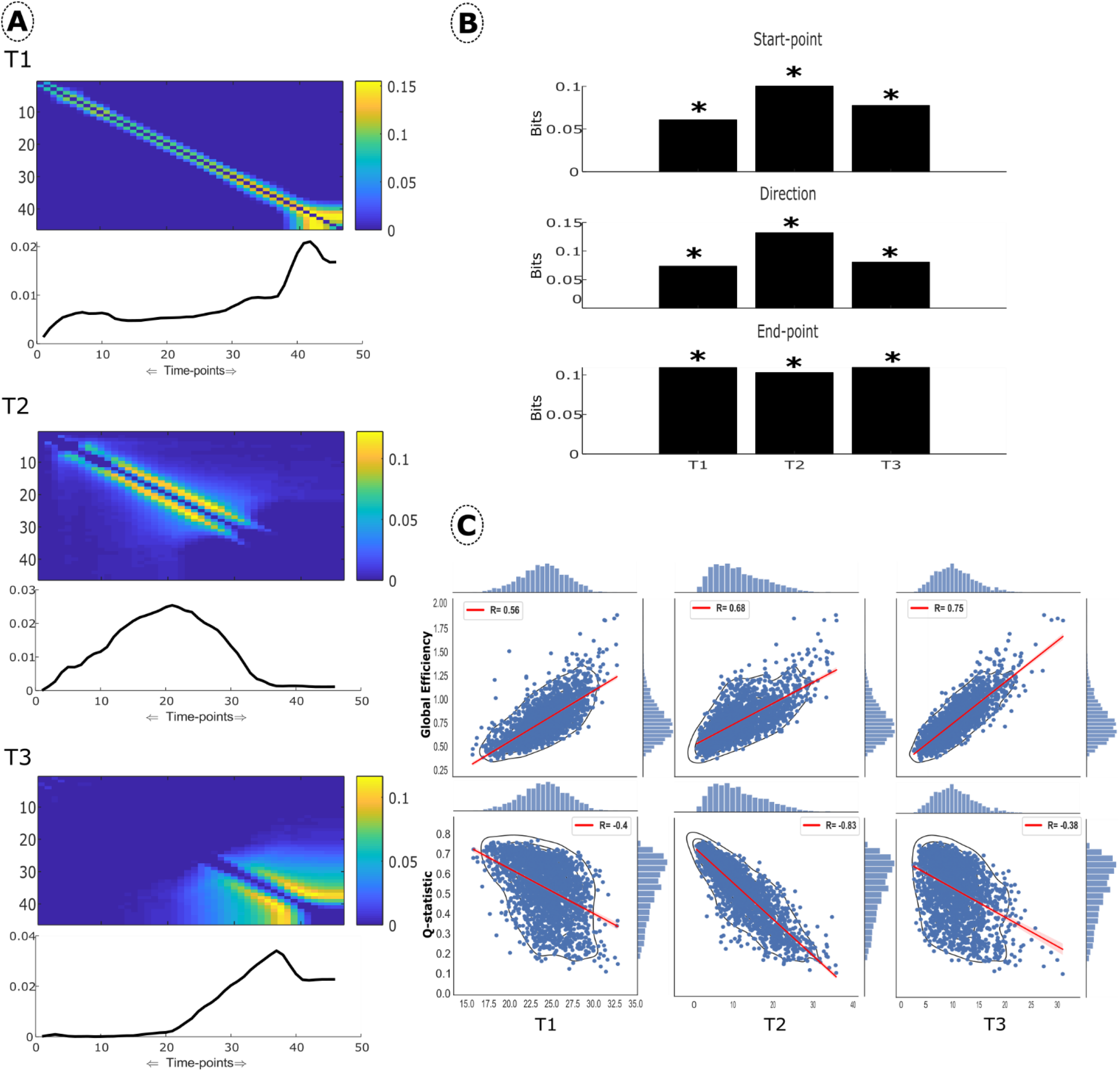
**(A)** Temporal synergies extracted from an example participant in dataset 2. Three communities were identified in the multiplex network and extracted using PNMF. The line plot below represents the average values within each column of the adjacency matrix. **(B)** The task-encoded information is presented in bits. Significant information (p<0.05) is indicated with *. **(C)** The noise correlations between each temporal synergy activation and Global Efficiency/Q-statistic. The bars along the axis of each plot are marginal histograms of the x- and y-variables.

#### Space-Time synergy model

We then sought to generalise the space-time synergy model results from dataset 1 to the example participant in dataset 2 (Fig.9). The findings were successfully replicated in a more complex dataset of whole-body reaching where a model-rank of three was found. ST1 here appears to capture the postural stabilisation exhibited predominantly in the upper-body but supported by the femoral muscles at movement onset and increasingly so at movement termination. ST2 is characterised by a phasic activation at movement initiation with a critical muscle (anterior deltoid) identified at the reaching shoulder supported by dependencies among the lower-limb muscles of both sides that counteract the induced shifts in centre-of-pressure. Following movement initiation, ST3 comes into a greater level of dependency between time-samples 12-28 approximately, accompanied by significant connections between the gluteals and among the right side lower-limb. This synergy then subsides back to its original activation level near movement termination, where the new body position is supported by the sustained activation in ST1. A similar level of dependency is found within the left upper-body and the right anterior deltoid in ST2 with this synergy reflecting the increased demands induced proximally in order to counteract the right arm during the transition to a new position.

**Fig.9:**
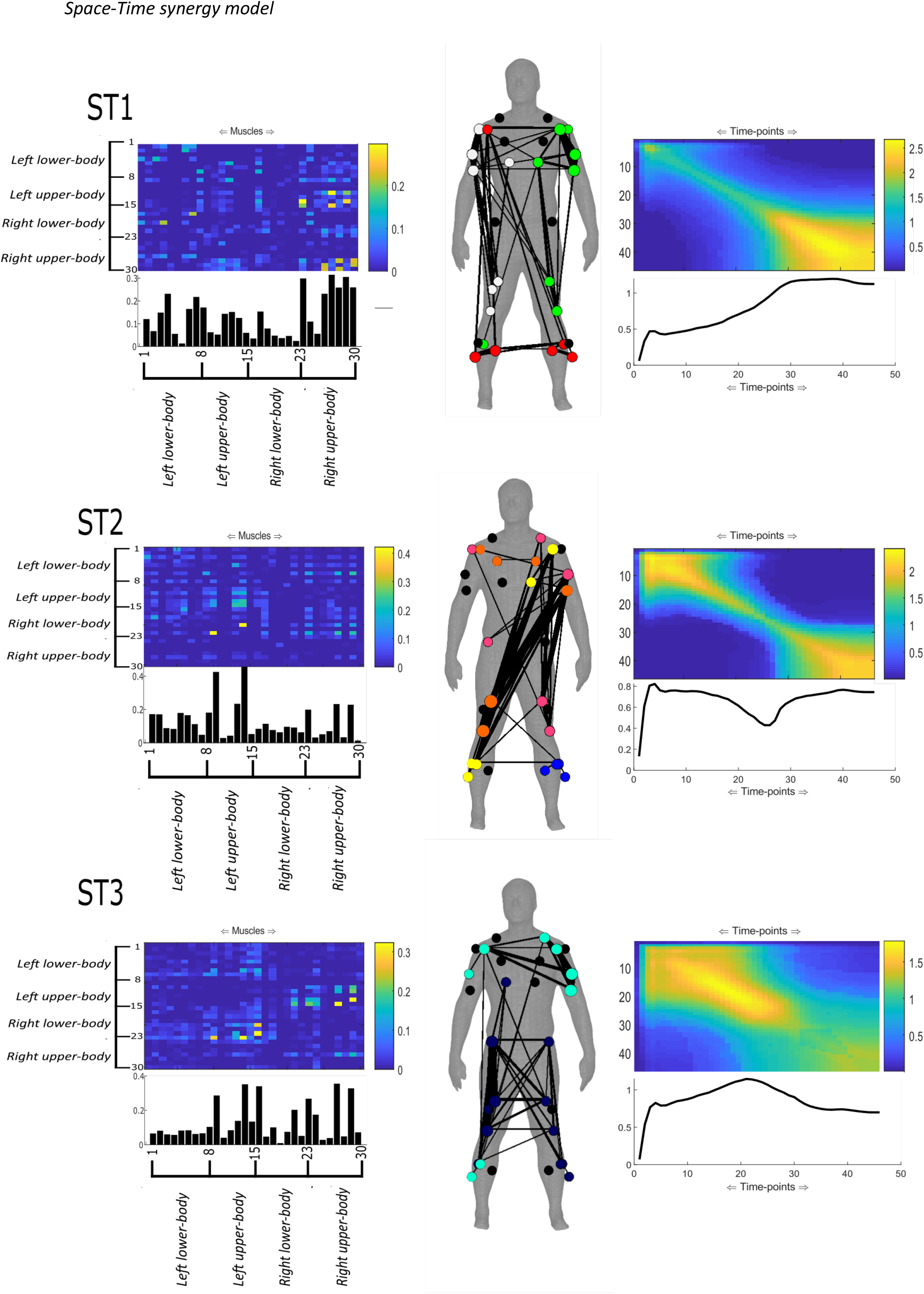
The Space-Time synergies extracted from the example participant in dataset 2. Three communities were identified and extracted using PNMF. Spatial and temporal synergies correspond on a 1:1 basis here as presented across rows. The spatial synergies are organised so that rows 1-8: left lower-limb, rows 9-15: left upper-body, rows 16-23: right lower-limbs and rows 24-30: right upper-body. The bar/line plots below represents the average values within each column of the corresponding adjacency matrix. The connection strengths, submodular structure and involvement of nodes are indicated by the minimally connected human body model via the edge widths, node colour and size respectively [57]. Submodular structure was identified using the conventional Louvain algorithm on the synergy matrices [47]. Unconnected nodes are in black.

A complex submodular structure was found across all three synergies, as exemplified on the human body models where three, five and two submodules were identified by a secondary community detection of ST1-ST3 respectively. The right anterior deltoid and tibialis musculature (red), right arm and femoral musculature (white) and left arm and femoral musculature (green) comprised ST1 submodules. ST2 consisted of strong connections between the left upper- and right-lower body (orange, yellow & wine) and a small cluster among left tibialis musculature (blue). ST3 comprised strong connections among the right lower-limb but also with the left leg (navy) along with a cluster of muscles across the upper-body that contained a long-range connection with the right soleus and anterior tibialis (cyan). Taking ST1 as an example here, the submodular architecture demonstrates the unique functional role of the reaching arms anterior deltoid which shares its community assignment with the lower-limb musculature. This submodular structure is likely representative of the reactive role the lower-limb musculature plays in reacting to stabilise the center-of-pressure against shifts in weight distribution of the reaching arm. The non-reaching arm comprises a submodule in both ST1 and ST3 that overlaps with the reaching arm anterior deltoid, indicating that despite being at rest, the non-reaching arm activations occur in synchrony across trials with the reaching arm and are indeed coupled to some extent.

#### Consistency of synergies (Dataset 2)

Following synergy extraction, we determined the consistency of identified spatial, temporal and space-time synergies across participants during whole-body reaching movements using a structural similarity index. Starting with the spatial synergy model, 3.4 (range=3-4) synergies were representative of the five participants in dataset 2 with the Q-statistic ranging from 0.997-0.998 and the threshold value 0.26 bits on average. The similarity index revealed a moderate level of correlation (R=0.56±0.19). Supplementary materials (Fig.5(A-C)) illustrates the average spatial synergies we extracted along with their respective task attribute dependencies and correlation with network properties. Despite this reduced similarity, the synergies are readily interpretable in their underlying functionality, likely indicating that the shape of activations rather than functionality was dissimilar across participants. For instance, S1 here consisted of dependencies within both the upper- and lower-body related to normal-stance stability and S2 is related to whole-body postural stability during reaching movements. As the greatest dependencies in S3 lie within the right upper-body, this synergy is representative of the activity in the reaching arm. These observations are supported by the task-encoded information. For instance for S2, the average task information was noticeably higher for start-point height (0.26 bits) and Up-down direction (0.28 bits) over other synergies, likely reflecting the differing postural adjustments required as the reaching arm is either raised up against gravity or down closer to a resting position. The trade-off between GE and Q captured in the activation coefficients was replicated across participants. Interestingly, fluctuations in S3 activations were most sensitive to changes in GE (R= 0.69) and Q (R= -0.57) across the remaining participants.

We then conducted this analysis on the temporal synergies of dataset 2 participants, producing a mean of 3.4 synergies (range=3-4) across participants that exhibited a high degree of concordance (R=0.91±0.57). This high level of concordance was accompanied by significant variability which we posit reflects the varying time-lags and contributions of individual muscles to reaching movements across participants that wouldn’t influence the overall shape of the synergy but the composition (i.e. higher/lower magnitude dependencies comparatively). The Q-statistic ranged from 0.97-0.995 while the average threshold value was 1.45 bits. In supplementary materials (Fig.6(A-C)), the representative temporal synergies for this sample are presented along with mean task-attribute and noise correlations with network properties. T2 was most highly modulated by start-point (0.05 bits) and direction (0.07 bits). T1 and T3 shared an equivalent level of modulation by end-point (0.546 and 0.547 bits respectively). The fluctuations in T3 activations were most sensitive to changes in GE (R= 0.69) and Q (R= -0.57), although all synergies demonstrated a high association across participants.

Finally, we applied the space-time model to all dataset 2 participants and their structural similarity was compared using a representative model rank of 2 (mean rank= 2.4, range=2-3). We found a satisfactory level of concordance across participants (R=0.79±0.4). Supplementary material (Fig.7) illustrates these representative synergies where the Q-statistic for modularity ranged from 0.93-0.997 and the threshold values ranged from 0.295-0.385 bits. ST1 functionally represents the deceleration and support of the reaching arm in its new position, which is reflected by the involvement of the upper- and lower-body of the opposing side. ST2 was active during the early to intermediary stages of the movement and mostly consists of couplings between lower-limb muscles, providing postural stabilisation during task execution.

## Discussion

To summarise the findings presented, functionally and physiologically meaningful muscle synergies were extracted using a network-information theoretic framework. Synergies were extracted in the spatial, temporal and spatiotemporal domains, each capturing unique sub-tasks and representing underlying mechanisms in human point-to-point reaching generalisable across two datasets that were consistent in their projections across participants. Along with this, we showed a significant task dependence of synergy activations that closely related to the functional interpretations of the extracted synergies and significant correlations with network properties that captured the dynamic trade-off between modularity and small-worldedness. Both model-rank selection and sparsification procedures were conducted prior to dimensionality reduction in a data-driven and holistic manner that reduces the necessity for post-hoc analyses. Through the determination of dependencies on a within-trial basis between muscles, time-sample vectors or their combination across trials, a novel formulation of muscle synergy was implemented that we suggest can provide important and novel insights into the neural control of human movement. In the following we detail some of the main commonalities of the GCMI framework with existing models along with novel findings and advantages. These commonalities and advantages are summarised in the Supplementary material of this article, where a formal comparison with existing models is provided (supplementary materials Fig.8).

### Continuity with previous muscle synergy models

Previous studies have been conducted on the datasets analysed here with interesting commonalities with our results [15,60,61]. Functional muscle groupings by the spatial synergy model here were reflective of previous findings (e.g. Dataset 1: elbow extensors (S1), shoulder flexors (S2) and shoulder and elbow extensors (S3); Dataset 2: a single synergy (S3) for the reaching arm along with whole-body functional groupings relevant for postural stability that cannot be explained by anatomical constraints). The spatial synergies presented in the current study were, similarly to previous reports, only moderately similar across participants in their shape but robust in their functional underpinning. This low level of agreement found in dataset 2 is likely related to physical discrepancies among participants (e.g. height, muscle length, motor preferences). The concordance among temporal synergies found in the current study also replicated previous work. Notably however, a high degree of inter-subject variability was reported here. As the temporal synergies consisted of dependencies between time-sample vector pairings across muscles, we posit that in contrast to the shape differences among spatial synergies, the overall shape of the temporal synergies was consistent, but their composition was not. The varying time-lags of individual muscles is likely to have contributed to this difference in composition [62,63], leading to comparatively higher/lower magnitude dependency at certain time-sample pairs across participants. This demonstrates the capacity for the GCMI framework to effectively capture the idiosyncrasies of individual participants’ movement patterns. To exemplify this point further, following a functional similarity analysis of temporal synergies it was noted in [61] that of the four synergies identified, several could be merged into fewer clusters. In the current study, these synergies were indeed merged into a readily identifiable bi-phasic pattern accompanied by an underlying tonic synergy, highlighting the functional relevancy of the synergies extracted here. The ability of GCMI to capture complex, nonlinear associations was advantageous in the merging of functionally synonymous phasic synergies (see ‘Materials and methods’ section) and exemplifies a step forward in the muscle synergy literature.

In terms of task dependence, significant shared information was observed in the activation coefficients of spatial and temporal synergies with various task attributes. An interesting difference in the task modulation of muscle synergies extracted here to that presented in previous work is that the synergies identified here convey information for several movement phases. This reflects the unique formulation introduced by the GCMI computations where early activations provide relevant task information about future states and vice-versa within the same vector in the input matrices. This allows for individual synergies to provide a more holistic insight into the task performed over existing approaches, representing synergies collectively as a single motor representation that unfolds across space and time. There was also some continuity with previous research. For example in dataset 2, whole-body reaching spatial synergies associated with postural stability were consistently modulated by task attributes that could be said to influence the participants medio-lateral or anterior-posterior centre-of-pressure (e.g. End-point height, Up-Down direction) as formerly observed [61]. The transient but also decelerative temporal synergy in dataset 2 (T2) was highly modulated by reaching direction, a replication of previous work also. The task modulation of space-time synergies has been reserved for investigation in future work.

### Novel insights from the GCMI framework

The GCMI framework presents a few advantageous qualities over existing approaches that have provided novel insights in the current study. Firstly, the interactions between pairs of muscles, timepoints or muscle-timepoint pairings is a novel characteristic of the GCMI framework. These pairwise interactions are uniquely present across all three GCMI models and reveal interesting (often overlapping) submodular structures representing the dynamics of muscle couplings at various temporal and spatial scales during movement. Furthermore, the use of GCMI to quantify these pairwise couplings reveals all muscle interactions (even the subtle ones) regardless of muscle amplitudes. This addresses a significant limitation of dimensionality reduction approaches to muscle synergy extraction which put more emphasis on muscles with a predominant role in the movement because these muscles explain most of the variance in the dataset.

In past work, the separation of tonic and phasic synergies has not been straightforward and commonly involved the subtraction of a linear ramp from EMG waveforms [64,65]. Within the GCMI framework, these distinct temporal synergy types are both isolated within the same extraction procedure (e.g. T1 in fig.5 and fig.8), leading to a data-driven approach that relaxes the assumption of linearity. In relating the mechanisms underlying the postural control of movement in the oculomotor system and the human arm, [66] elucidated a plausible mechanism for subcortical postural control. The integration of cortically generated movement commands by a separate, subcortical postural controller was initially supported in primates and found to be generalizable to healthy human and cortically-impaired populations where this dependence persisted on a within-trial basis only. In the current study, tonic synergies were identified in reaching tasks on a within-trial basis only that consistently presented a pattern of activation indicative of the aforementioned findings. More specifically, a low-to-moderate gradually increasing/decreasing level of dependency was found throughout T1 (fig.5 and fig.8) until a sharp spike in dependency was found consistently across participants at the last few time-sample pairings, potentially representing the integration of the preceding movement activations in the holding response. Further analysis using higher resolution EMG data along with relaxations in rigid temporal alignments may provide further insight into the underlying mechanisms governing motor control. This increased insight is made possible by key advantages provided by this approach in terms of flexibility and multiplexity with findings that may run in parallel to recent innovations in time-warped tensor decompositions [67].

The ability for a self-organising system to re-organise its elemental variables from trial-to-trial to complete a given task is an important attribute of the synergy concept [29]. In the development of this framework, careful consideration for important network properties characteristic of biological systems was made throughout the analysis. We chose to focus on two opposing but essential network properties, fractal modularity and small-worldedness, due to their functional implications and focus in the relevant literature [41,43,49–51,68]. The preservation of these important network properties in the extracted synergies was confirmed through the identification of significant noise correlations in the underlying activation coefficients with trial-to-trial fluctuations in modularity and global efficiency. The persistent opposing directions and strength of these associations with GE and Q are both reflections of this trade-off and the sensitivity of the extracted synergies to trial-to-trial fluctuations at the network level respectively. These findings are also in line with the current understanding of the phenomena of adaptive breakdowns in modularity in complex systems and findings relating gain modulation to shifts in network topology [68,69]. Furthermore, the maintenance of fractal modularity in the output of the GCMI framework is exemplified by the consistent presence of functionally meaningful submodules within the extracted synergies. It is likely that many of these submodules were defined by biomechanical constraints and a localised function [23,24], however, a significant number of long-range connections clearly transcended this constraint (e.g. the right anterior deltoid and lower-limb musculature in the human body models of dataset 2). It is also noteworthy that many of the identified submodules overlapped, indicating that many individual muscles demonstrate a complex functionality that is shared across a distributed subnetwork of muscle couplings, supporting previous findings [23–25]. We thus suggest that the synergies extracted through the GCMI framework may be useful for investigating the unique contribution of both biomechanical and neural constraints on movement.

The identification of movement phase commencement and cessation has been frequently cited in the literature as difficult, especially during fast-paced movements due to overlapping muscle activations [65,70]. The orthogonality introduced by PNMF here effectively removes this limitation by minimising the overlap between temporal synergies. The trial-specific activation coefficients produced by the proposed spatial and temporal models here significantly improve the analytical flexibility and convenience of the current models which are difficult to contrast against other variables of interest. The framework is not only convenient for further analysis but can provide novel fundamental insights. For example, in both datasets a novel bi-phasic pattern supported by an underlying tonic synergy was revealed. This bi-phasic pattern is reflective of the bang-bang control policy investigated recently that was found to explain the characteristic tri-phasic activation of reaching movements [70]. The findings presented here support the hypothesis of an intermittent rather than continuous control scheme in human motor control and implicate this control policy as a fundamental mechanism underlying human motor control [71,72].

Muscle synergy analysis is known to be scale-dependent, in that the number and spread of muscles analysed across the body influences the synergy output, typically by increasing model-rank with increasing dataset complexity [45]. Instead of optimising the variance accounted for criterion, the proposed framework identified a reduced number of spatial and temporal synergies that were invariant to dataset complexity by optimising a more task-relevant modularity criterion. This is a significant progression from existing approaches which rely solely on variance optimisation frequently at the expense of task relevancy [22]. Within the sensory system the multiplex encoding of information across multiple time-scales is well-known [73], allowing for an increased coding capacity and the apparent continuous stream of perception. This multiplexity has more recently been identified in the encoding of task information from low-dimensional dynamics in a motor cortical neural population and put forward as a structural mechanism underlying cognition and self-organisation more generally [74–76]. Other recent research that has modelled muscle synergies as a multiplex network only identified a single module representing a global co-activation during gait in various coordination modes [26]. This global co-activation may have masked more discrete but important mechanisms that could not be uncovered by simpler methodologies. In the current study, the multiplexity of human motor control was exhibited through the parsing of several community structures across multiple spatial and temporal scales that demonstrated significant encoding of task information. Therefore, we suggest that the GCMI framework presents as a useful tool for investigating the emergence of complex patterns in motor behaviour.

### Future research directions

A key motivation for the development of this framework was to address the limitations of linear assumptions and lack of flexibility in current muscle synergy models. The inclusion of task space variables has recently come into the spotlight in this line of research [11], as the pattern configurations found are required to be constrained by an objective function in order to be functionally relevant and transferable to robotics and prosthetic design [11,22]. Through the induction of non-parametric statistical tools in information and network theory, the presented formulations are now more amenable to the inclusion of task space variables while more appropriately capturing the non-linearity of the musculoskeletal system. The objective function to constrain these extracted synergies may also involve the inclusion of neural data derived elsewhere in the CNS, producing neurophysiological relevant synergies that are falsifiable [1,5]. Each muscle synergy model may capture distinct motor features [77], thus investigating their functional underpinning following the inclusion of these constraints may be fruitful. Information theory more generally has found great use in the analysis of complex systems, for instance in identifying synergistic and redundant interactions in the CNS, the integration of cross-modal informational dynamics and providing a framework for understanding causal emergence [78–80]. Quantifying informational dynamics among muscles in space and time may allow for the identification of task-relevant and irrelevant spaces, providing a bridge between neurophysiological and purely computational frameworks [81,82].

