## Supplementary materials for "A network information theoretic framework to characterise muscle synergies in space and time"

**Supplementary Material**

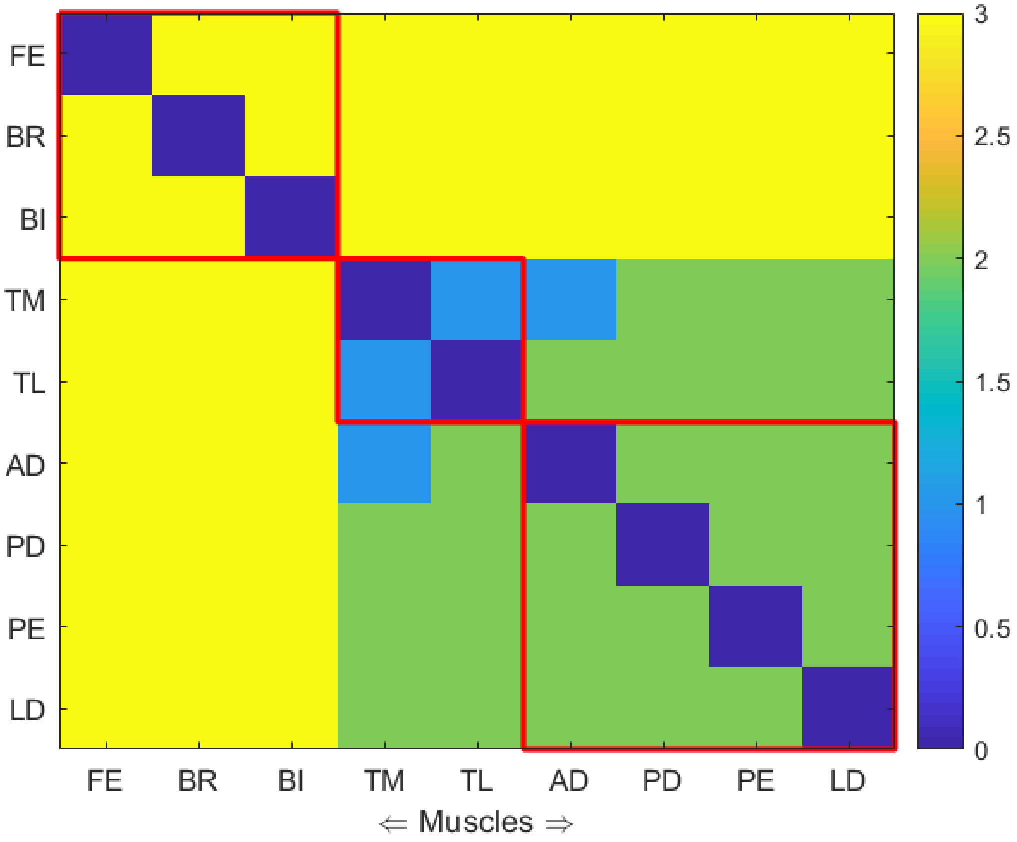


**Fig.1**: An example of the cluster assignment verification procedure described in the Materials and methods section applied to dataset 1. The red grid lines represent the hard-clustering assignment provided by the generalised Louvain algorithm while the colours in the adjacency matrix indicate the cluster assignment produced by PNMF.

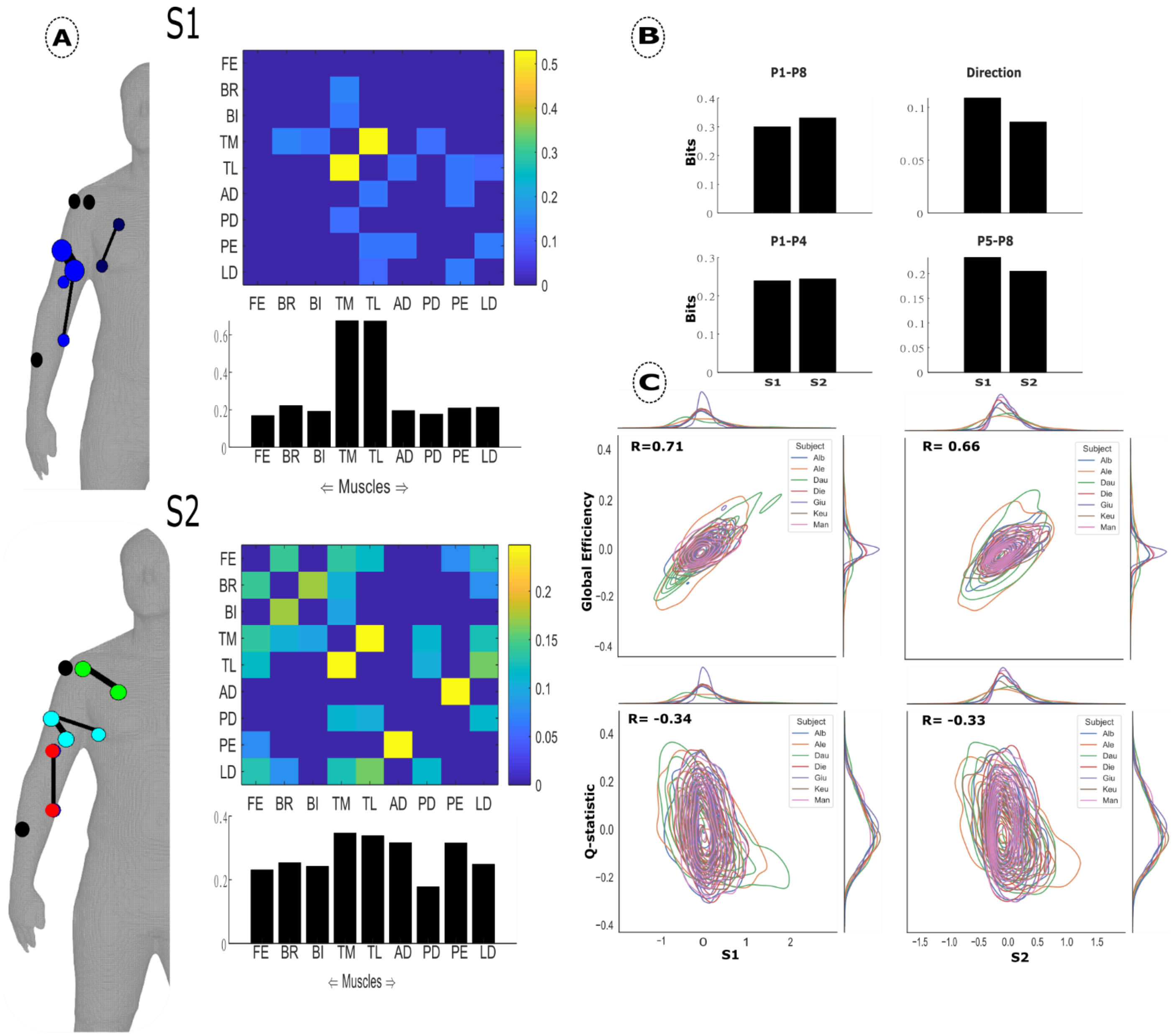


**Fig.2**: **(A)** The average spatial synergies across participants. The bar graph below represents the average values in each column of the adjacency matrix. The human body models illustrates the values in the adjacency matrix with the width of the edges and colour and size of the nodes providing insight into connection strengths, modular structure and involvement respectively [57]. Submodular structure was identified using the conventional Louvain algorithm on the synergy matrices [47]. Unconnected nodes are in black. **(B)** The average task encoded information for two spatial synergies found to be representative of dataset 1 participants. The task attributes analysed included: P1-P8, P1-P4, P5-P8 and Direction. **(C)** Average correlation between trial-to-trial fluctuations in Global Efficiency/Q-statistic for modularity and noise in the spatial activation coefficients across participants. The curves along the axes of each plot are kernel density estimates for each participants' x- and y-variables.

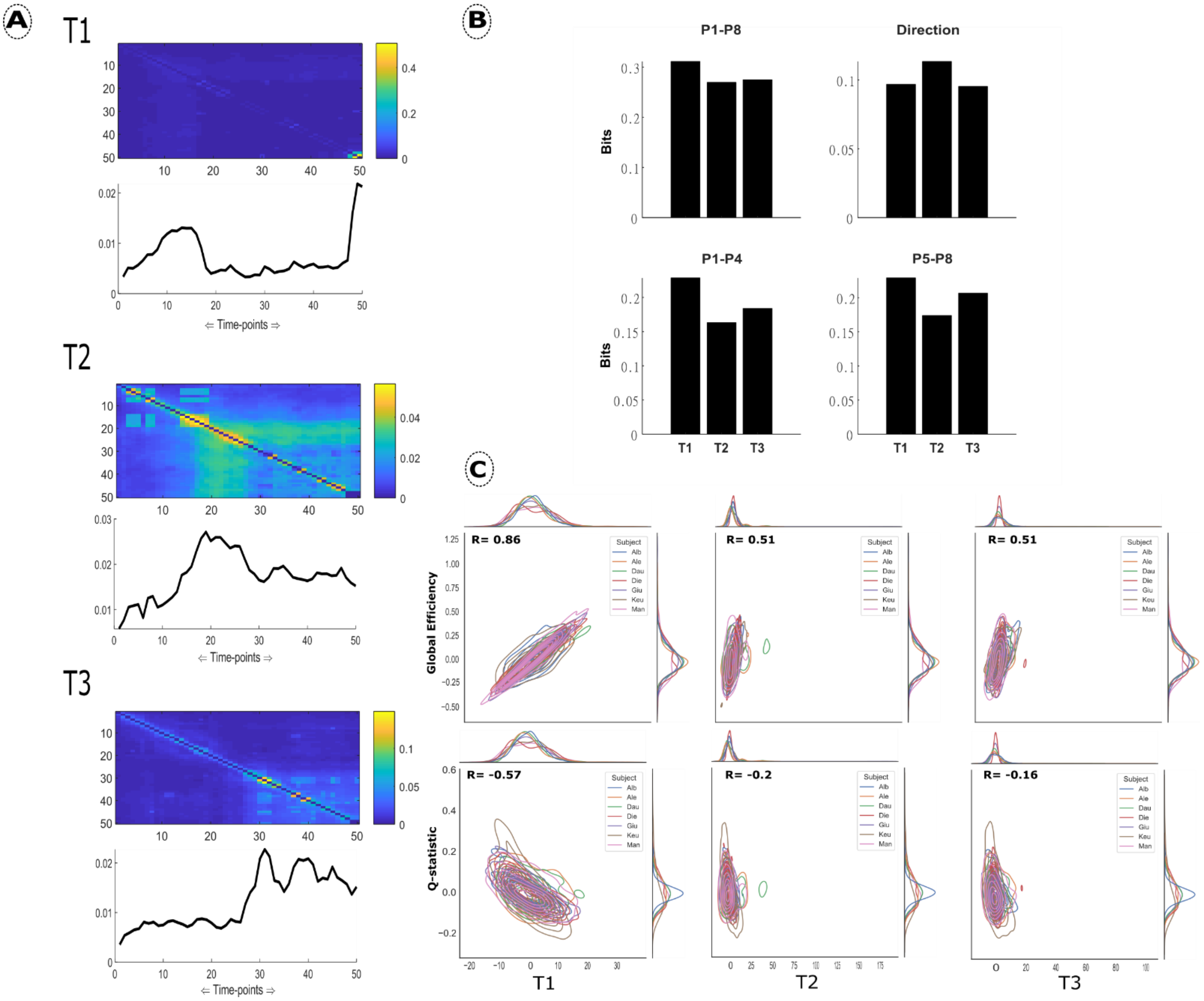


**Fig.3**: **(A)** The average temporal synergies across participants in Dataset 1. The line plots represent the average of each column in the adjacency matrix above. **(B)** The average task encoded information for three temporal synergies found to be representative of dataset 1 participants. The task attributes analysed included: P1-P8, P1-P4, P5-P8 and Direction. **(C)** The average noise correlation between temporal synergy activations and Global Efficiency/Q-statistic across participants. The curves along the axes of each plot are kernel density estimates for each participants' x- and y-variables.

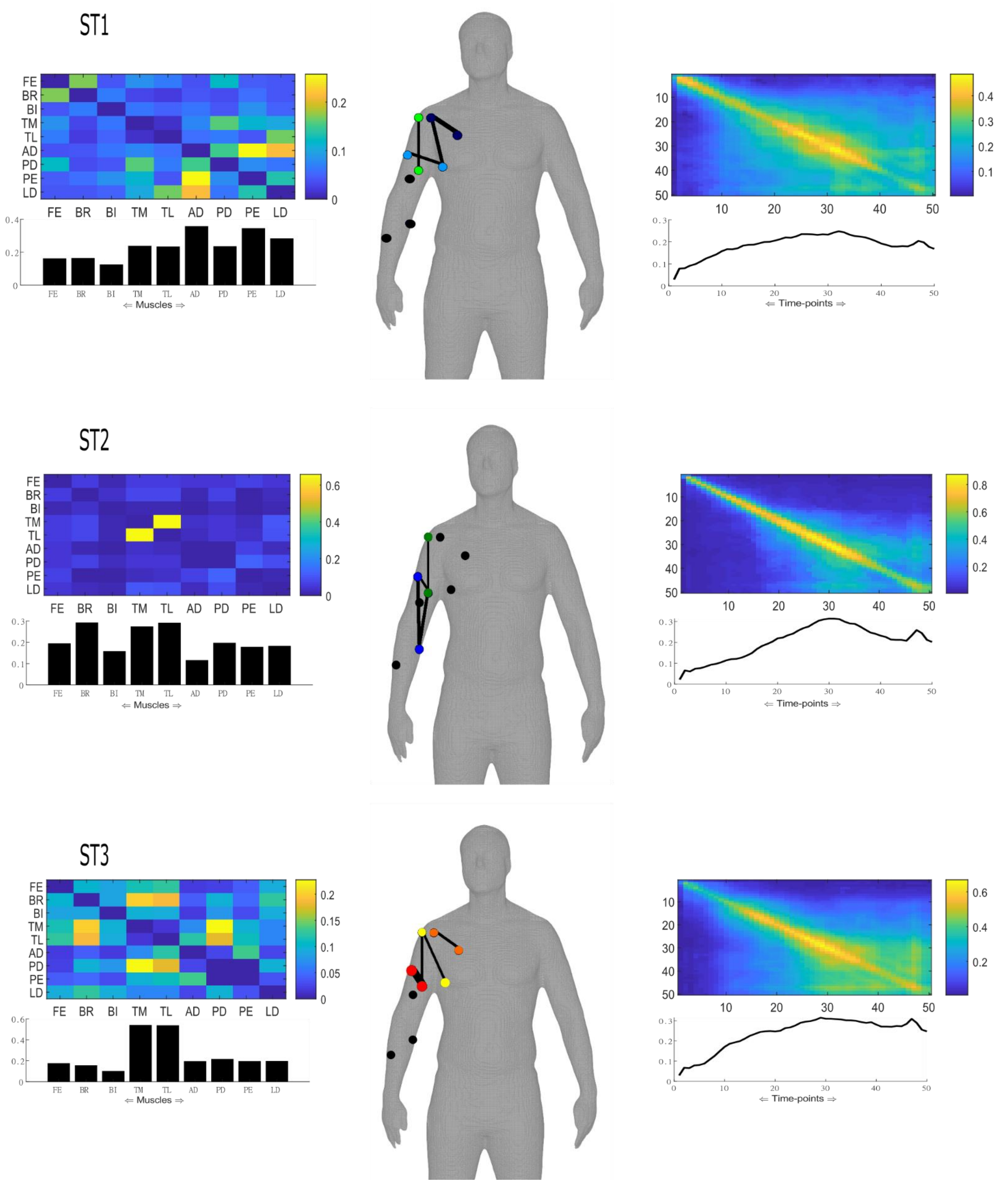


**Fig.4:** The average space-time synergies across participants in dataset 1. Spatial and temporal synergies correspond on a 1:1 basis as shown across rows. The bar/line plots represent the average of each column in the adjacency matrix above. The human body model illustrates the values in the adjacency matrix with the width of the edges and colour and size of the nodes providing insight into connection strengths, submodular structure and involvement respectively [57]. Submodular structure was identified using the conventional Louvain algorithm on the synergy matrices [47]. Unconnected nodes are in black.

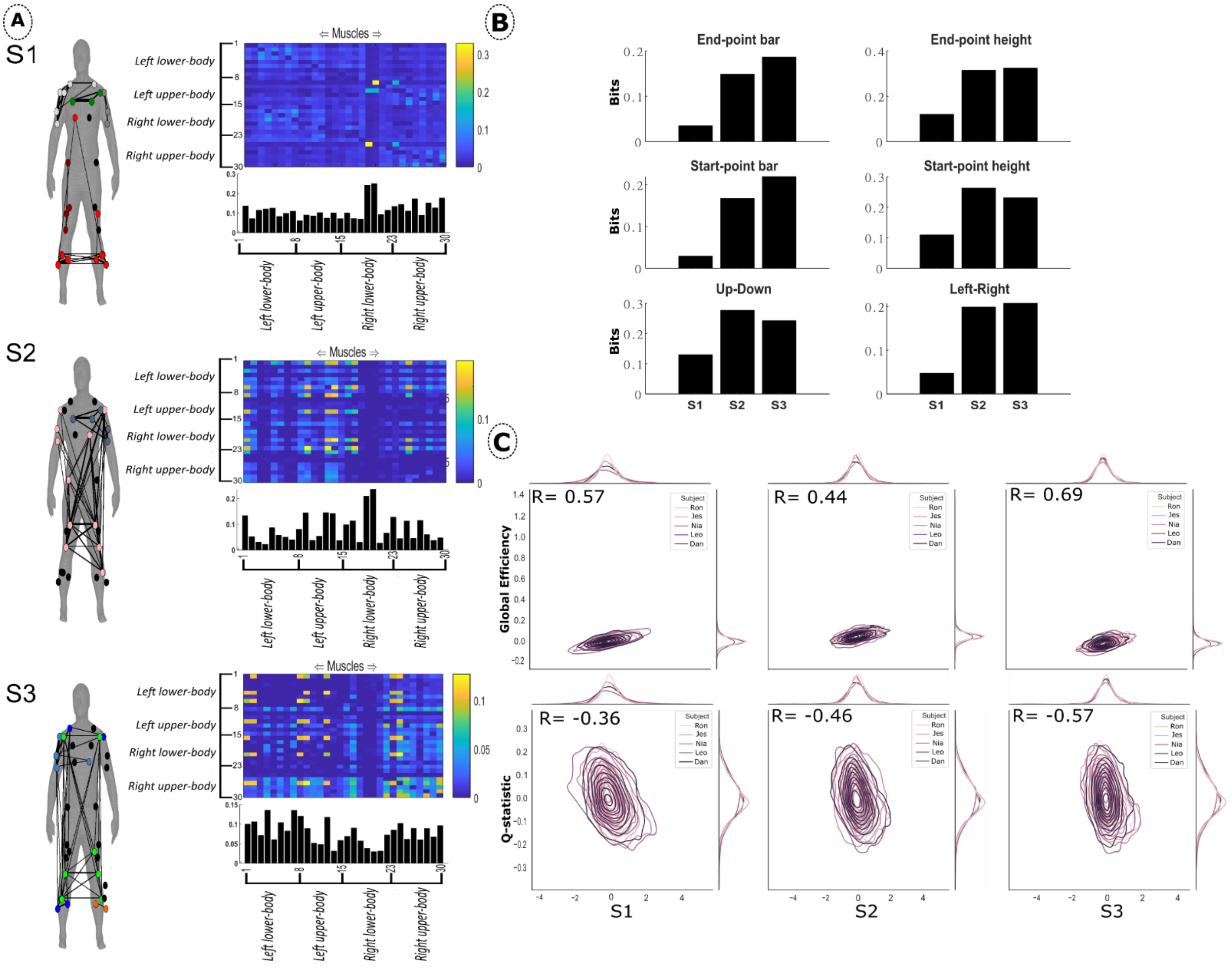


**Fig.5**: **(A)** The average spatial synergies across participants in dataset 2. The adjacency matrices are organised so that rows 1-8: left lower-limb, rows 9-15: left upper-body, rows 16-23: right lower-limbs and rows 24-30: right upper-body. The width of the edges on the human body model and the size and colour of the nodes indicate the connection strength, node involvement and submodular structure respectively [57]. Submodular structure was identified using the conventional Louvain algorithm on the synergy matrices [47]. Unconnected nodes are in black. **(B)** The average information encoded for six task attributes is presented in bits. **(C)** The average noise correlations between spatial synergy activations and Global Efficiency/Q-statistic across participants. The curves along the axes of each plot are kernel density estimates for each participants' x- and y-variables.

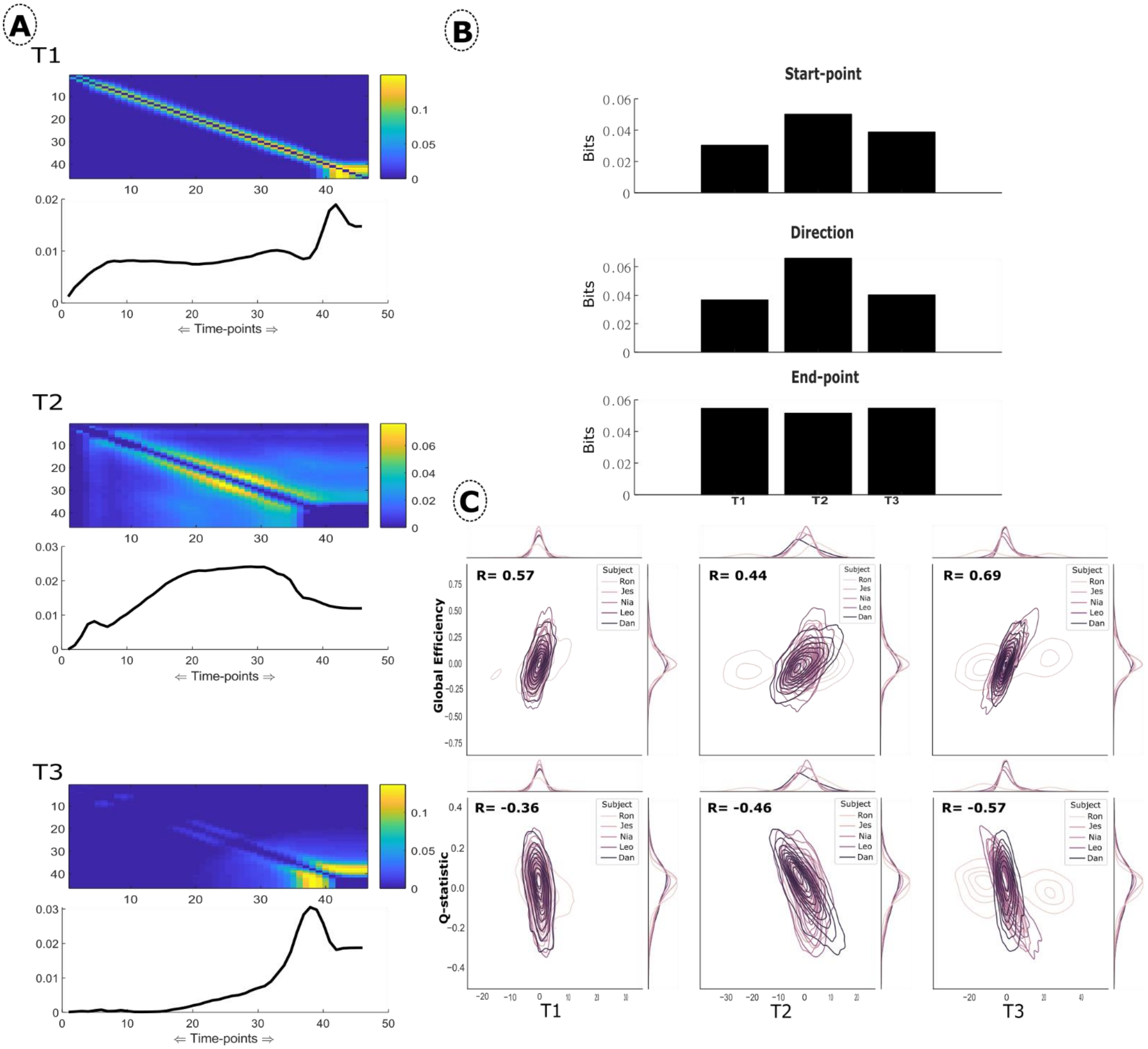


**Fig.6**: **(A)** The average temporal synergies extracted from dataset 2. The line plots represent the average of each column in the adjacency matrix above. **(B)** The mean task-encoded information for start- and end-points and direction. **(C)** The average noise correlations between temporal synergy activations and Global Efficiency/Q-statistic across participants. The curves along the axes of each plot are kernel density estimates for each participants' x- and y-variables.

ST1

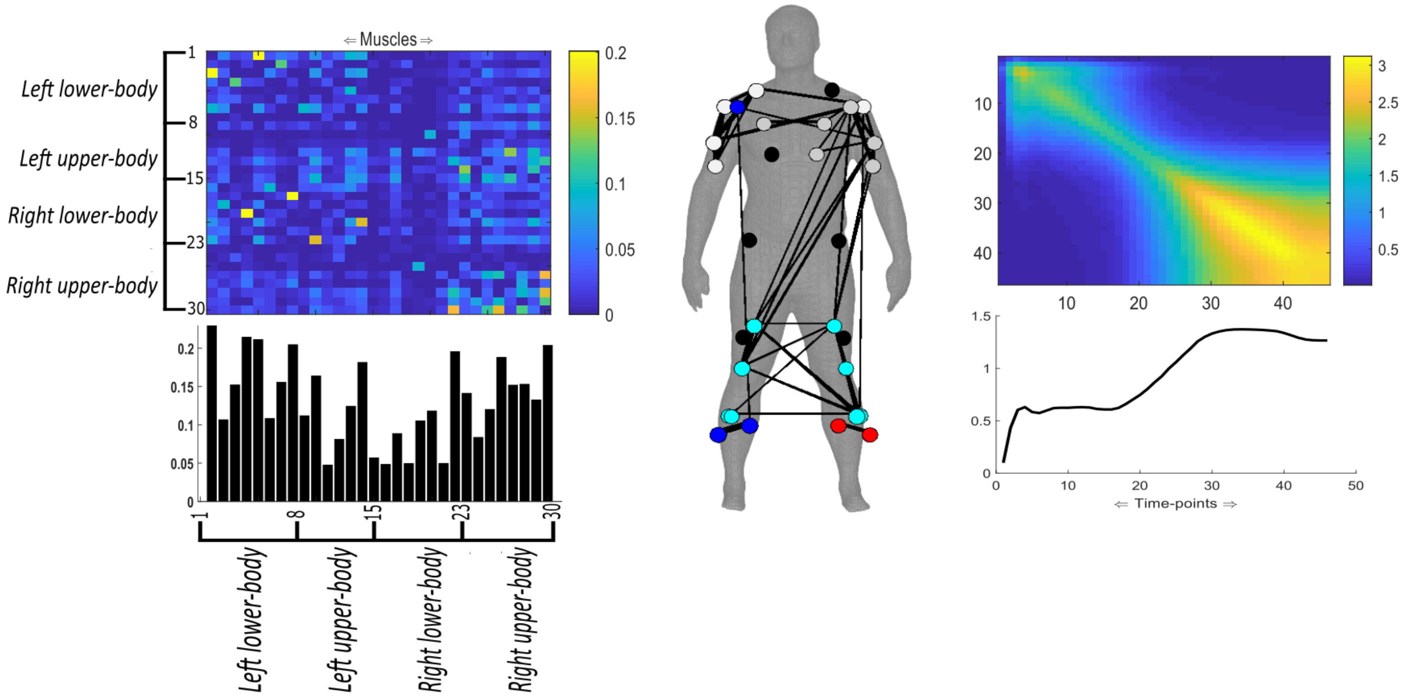


ST2

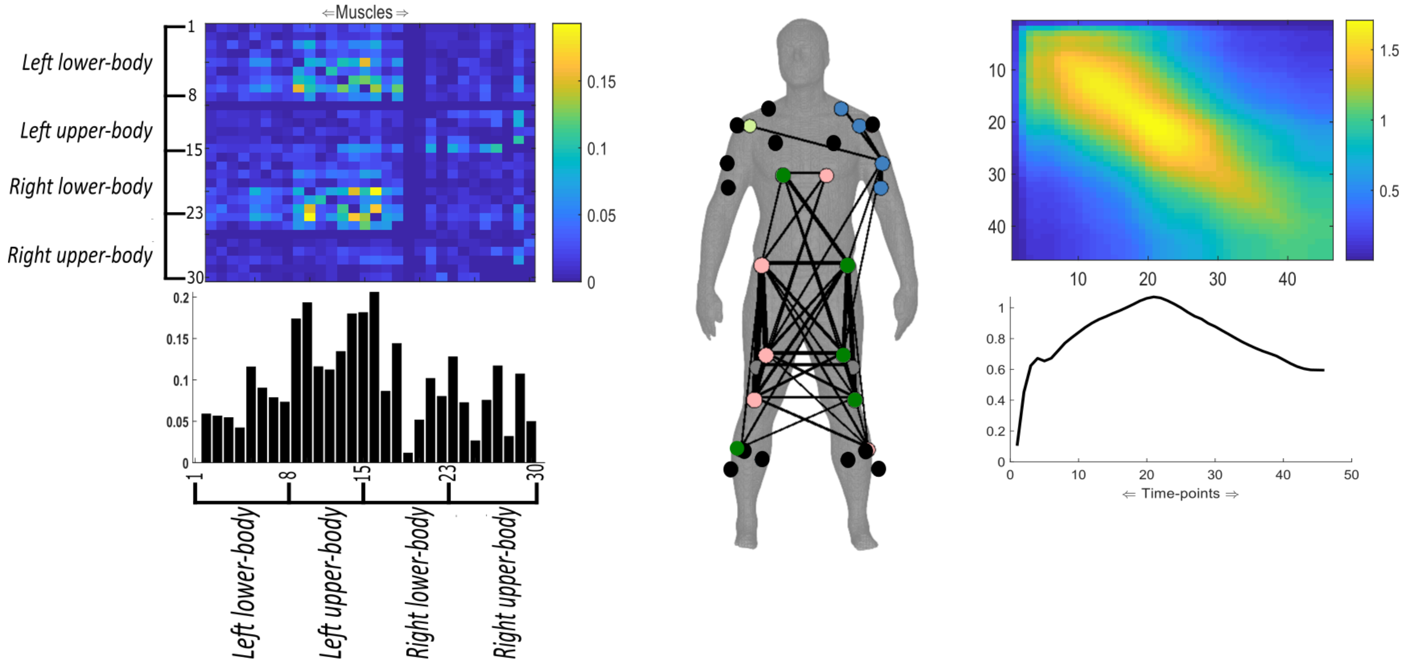


**Fig.7**: The average Space-Time synergies from dataset 2. Spatial and temporal synergies correspond on a 1:1 basis here as presented across rows. The spatial synergies are organised so that rows 1-8: left lower-limb, rows 9-15: left upper-body , rows 16-23: right lower-limbs and rows 24-30: right upper-body. The bar/line plots represent the average of each column in the adjacency matrix above. The connection strengths, submodular structure and involvement of nodes are indicated by the human body model via the edge widths, node colour and size respectively [57]. Submodular structure was identified using the conventional Louvain algorithm on the synergy matrices [47]. Unconnected nodes are in black.

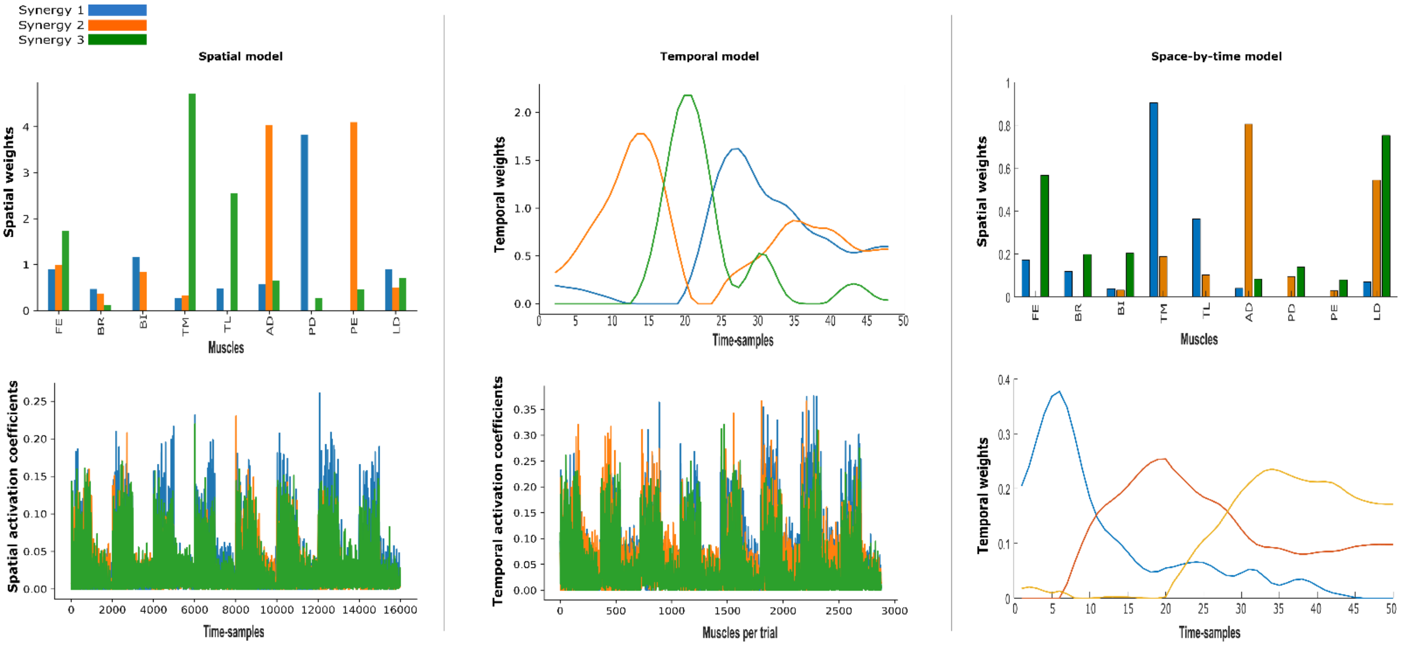


**Fig.8:** The application of current muscle synergy models to the example participant in dataset 1. A model rank of three was selected for all to assist in comparability with the GCMI frameworks output. Both the spatial and temporal models' synergies were extracted using non-negative matrix factorisation. The synergies presented for the space-by-time model were extracted using a non-negative matrix tri-factorisation method [15].

### Comparison of proposed framework with existing approaches

An aim of this study was to develop a framework for the extraction of muscle synergies that would alleviate limitations in the current models including linearity and a lack of flexibility and generalisability. Here, we wish to further highlight some of the commonalities of proposed GCMI framework with existing models and outline the advantages of the proposed framework. In Fig.8 of this document, we present the output from an application of the spatial, temporal and space-by-time models to the nine muscle EMG data of the same example participant (Dataset 1) presented in the main text. This dataset consisted of 320 trials of fast reaching movements in various targets and in forward and backward directions.

Starting with the spatial synergy model, the medial- and lateral triceps (TM,TL) are predominant in the third synergy while more proximal musculature surrounding the shoulder joint (anterior deltoid (AD) and pectoralis major (PE)) are emphasised in the second synergy (Fig. 8). This grouping of muscles with respect to anatomical proximity is also captured in the GCMI spatial synergy model (Fig.5 of main text). However, the coupling between the finger extensors (FE), brachioradialis (BR) and biceps brachii (BI) is uniquely captured in S3 of Fig.5 (main text) while the only feature these muscles share in the spatial model of Fig. 8 is a shared relatively low-weighting across all three synergies. This example demonstrates how the current spatial synergy model, due to its reliance on variance explained, may put an emphasis on muscles with a predominant role in the movement and which explain most of the variance in the dataset. This inevitably comes at the expense of more subtle couplings such as in the forearm musculature here.

The temporal synergies produced by the GCMI temporal model are highly consistent with what is produced by the current temporal and space-by-time models. In contrast, the output of the space-time model we introduced here differs significantly from any other model, with a co-activation synergy and two tonic waveforms. This is probably a result of the current formulation of the space-time model, which simultaneously extracts spatial and temporal modules that are consistent across trials. This trial-by-trial consistency is a unique feature of the space-time model that can be used to reveal couplings that are shared across tasks or experimental conditions. For instance, ST3 is unique to the GCMI output with no such shared weighting among the triceps with the posterior deltoid (PD) found in the space-by-time model output.

Another notable advantage of the GCMI formulation is that the specific timing of movement onset and cessation is more interpretable due to the orthogonality introduced during cluster extraction. For example, T2 of Fig.6 (main text) illustrates a phasic burst of activation at time-sample 15-23 approximately that share information with the remaining time-samples in the movement, an attribute not present in the current muscle synergy models.

More importantly, the interaction between individual muscle, time-sample or muscle - time-sample pairings is a novel characteristic of the GCMI framework. As recent evidence has shown [23-25], a single muscle may belong simultaneously to a number of functional groups, pertaining to the various functional roles a muscle can take on for a given task (e.g. internal joint stiffness, acceleration-deceleration, postural stabilisation etc.). These pairwise interactions are uniquely present across all three GCMI models, revealing submodular structures that may represent these complex interactions and cannot be inferred by the traditional synergy extraction approaches. The reliance of the current models on dimensionality reduction furthers the significance of this point as the more subtle interactions are potentially not captured at all.

Moreover, the model-rank procedures implemented by the existing approaches seek to optimise the variance accounted for. Although many important findings have been produced, this approach may limit the generalisability and task relevancy of the synergy output [11,22]. Within the GCMI framework, we implement a data-driven approach for model-rank selection that relies purely on the data structure and the number of identifiable clusters a priori, thus alleviating this limitation.

Finally, the activation coefficients, which are inferred to be the neural commands driving the muscle patterns, are captured across time-samples and across muscles per trial for the spatial and temporal models respectively (supplementary materials Fig.8). This output is difficult to contrast against any specific trial or task, a difficulty overcome by the space-by-time model. In contrast, the GCMI formulation of the spatial and temporal models produces trial-specific modulations which can be easily assessed against other variables of interest, offering greater flexibility in analysis but also a unique within-trial view into the underlying spatial or temporal dynamics.
